# Orbitofrontal cortex populations are differentially recruited to support actions

**DOI:** 10.1101/2022.08.08.503227

**Authors:** Christian Cazares, Drew C. Schreiner, Mariela Lopez Valencia, Christina M. Gremel

**Author notes:** Corresponding Author: Christina M. Gremel, Ph.D. University of California San Diego, 9500 Gilman Drive, Mail Code 0109 La Jolla, CA 92093-0109.

## Abstract

The ability to use information from one’s prior actions is necessary for decision-making. While Orbitofrontal cortex (OFC) has been hypothesized as key for inferences made using cue and value-related information, whether OFC populations contribute to the use of information from volitional actions to guide behavior is not clear. Here, we used a self-paced lever-press hold down task in which mice infer prior lever press durations to guide subsequent action performance. We show that activity of genetically identified lateral OFC subpopulations differentially instantiate current and prior action information during ongoing action execution. Transient state-dependent lOFC circuit disruptions of specified subpopulations reduced the encoding of ongoing press durations but did not disrupt the use of prior action information to guide future action performance. In contrast, a chronic functional loss of lOFC circuit activity resulted in increased reliance on recently executed lever press durations and impaired contingency reversal, suggesting the recruitment of compensatory mechanisms that resulted in repetitive action control. Our results identify a novel role for lOFC in the integration of action information to guide adaptive behavior.

## Introduction

Flexible decision-making requires successful use of information derived from past experiences.^1–4^ Orbitofrontal cortex (OFC) has been hypothesized to process inferred information relevant to ongoing task demands, integrating inferences into a “cognitive map” to support ongoing decision-making processes.^5–11^ Past investigations have supported this hypothesis, showing OFC activity contributes to inferred information derived from externally-derived sources, such as with predictive cues that can elicit behavior,^12–16^ cued choices,^17–22^ and outcome value.^23–30^ However, prior actions can also be used as information for inferences critical to adaptive control^3, 31–33^ and whether OFC populations are recruited for use of such action-related information has been debated.^34^ This is important to resolve as OFC activity is disrupted across psychiatric disorders involving aberrant action control.

Volitional actions provide one the ability to dictate opportunities and achieve desired outcomes.^35^ This differs from situations where behavior can be elicited or signaled by external sources in the environment. Instead, self-generated actions are organized and initiated based on inferences that arise within one’s self.^36^ For example, a road sign can signal which way to walk to get ice cream, or one can infer from prior experiences which direction to go. Depending on recent experiences, one can repeat actions to exploit a known rule, or modify an action to explore for new rules,^37, 38^ allowing one to adjust behavior from one decision to the next. However, whether such action-related inferences recruit OFC-based contributions is less clear. On one hand, prior investigations have observed modulation of lateral OFC (lOFC) neurons during actions^39–42^ and found that disrupting lOFC activity perturbs actions sensitive to decreases in expected outcome value.^40, 43–46^ Additional work has suggested lOFC is recruited when action-outcome contingencies change during learning.^47^ In contrast, studies have suggested that OFC populations may not participate in action processes per se, but instead are only recruited when cue-related, outcome-related, or value-related information is also present.^13, 48, 49–55^ In support of the latter hypothesis, broad (i.e. not population specific) chemical-induced lOFC inactivation in marmosets was found to enhance choice sensitivity to changes in action-outcome contingencies, raising the possibility that OFC performs functions that compete with action control processes.^56^ However, procedures often used to assess or degrade action-outcome contingencies or to decrease the value of associated outcomes do not provide a way to examine adjustments to the action itself, independently from adjustments to the relationship between an action and its associated outcome. Furthermore, relatively longer-term (i.e. lesions or whole-session manipulations) and non-specific lOFC activity disruptions found in these aforementioned studies may have facilitated compensatory mechanisms that could assume responsibility for the observed behavioral disparities.^44, 57, 58–60^ Thus, the specific contributions of lOFC to action-related information, if any, remain ambiguous.

Lateral OFC is widely innervated by cortical, thalamic, and subcortical areas,^61–66^ with incoming afferents synapsing onto various cortical cell types that include excitatory projection neurons and local interneuron populations that shape local network rhythmicity and neuronal firing.^67–71^ Despite this vast interconnectivity, little is known about how information used for inferences is integrated within these lOFC microcircuits. As different cell-types may receive similar inputs, and thus potentially similar information, genetically distinct lOFC subpopulations could show functional homogeneity or differential representation of information used for inferences that guide adaptive behavior.

Here we investigated whether lOFC projection and local inhibitory populations are important for volitional action control. We used a self-paced instrumental task in which prior actions and the continuous context in which they occur are used to guide subsequent actions.^33, 42^ In this task, mice had to learn to press and hold a lever down beyond a fixed minimum duration criterion to earn a food reward, with the reward only delivered after a successful lever press was terminated. This allowed us to examine adjustments to action control while keeping broad action-outcome relationships stable. Mice learned to adjust lever press durations (i.e. an analog measure) to earn rewards solely based on prior experiences, with no external cues or predictive information available to guide their ongoing behavior. Furthermore, lever pressing was goal-directed in that it was sensitive to shifts in action contingencies and decreases in expected outcome value. We performed calcium-based fiber photometry of genetically distinct lOFC populations and found representation of volitional actions and action history in lOFC Calcium/calmodulin-dependent protein kinase II (CamKII+) projection neurons, and to a much lesser degree, in Parvalbumin (PV+) inhibitory interneurons. Behavior-dependent optogenetic perturbations to distinct lOFC populations showed that lOFC activity encoded action-related information but was not critical for using prior action information to guide performance. Functional chronic loss of lOFC circuits produced a greater reliance on the most recently executed action despite a change in action contingencies. This suggests that a loss of lOFC resulted in recruitment of compensatory mechanisms and circuits that produced repetitive action control at the expense of integrating actions with broader experiential information for behavioral control. As such, we hypothesize that lOFC performs computations that contribute to the use of action history to guide adaptive behaviors.

## Results

### Mice learned to adjust self-generated lever presses using inferred action-related experience

Behavior is shaped in real time by its history. We adapted a lever-press hold down task in which mice had to learn to hold down a lever press for a duration longer than an arbitrary, unsignaled and predetermined minimum amount of time. Importantly, reward delivery only occurred immediately after the termination of any lever press that successfully exceeded this duration criterion.^33, 42, 72–75^ Thus mice have only their experience, including prior lever press durations, to guide subsequent action performance. Briefly, mice (C57BL/6J n = 23, 13 males, 10 females; PV^cre^ n = 14, 11 males, 3 females; no effect or interaction of genotype or sex on any behavior measure, thus groups were combined in subsequent analyses) learned to hold down a lever press for longer than an arbitrary duration to earn a reward (Figure 1A). Lever pressing was self-initiated, self-paced, and the lever remained extended into the chamber for the entire session. Importantly, there were no cues predictive of reward and mice received no feedback of performance success or failure until they terminated the lever press with reward delivered at offset.

**Figure 1.**
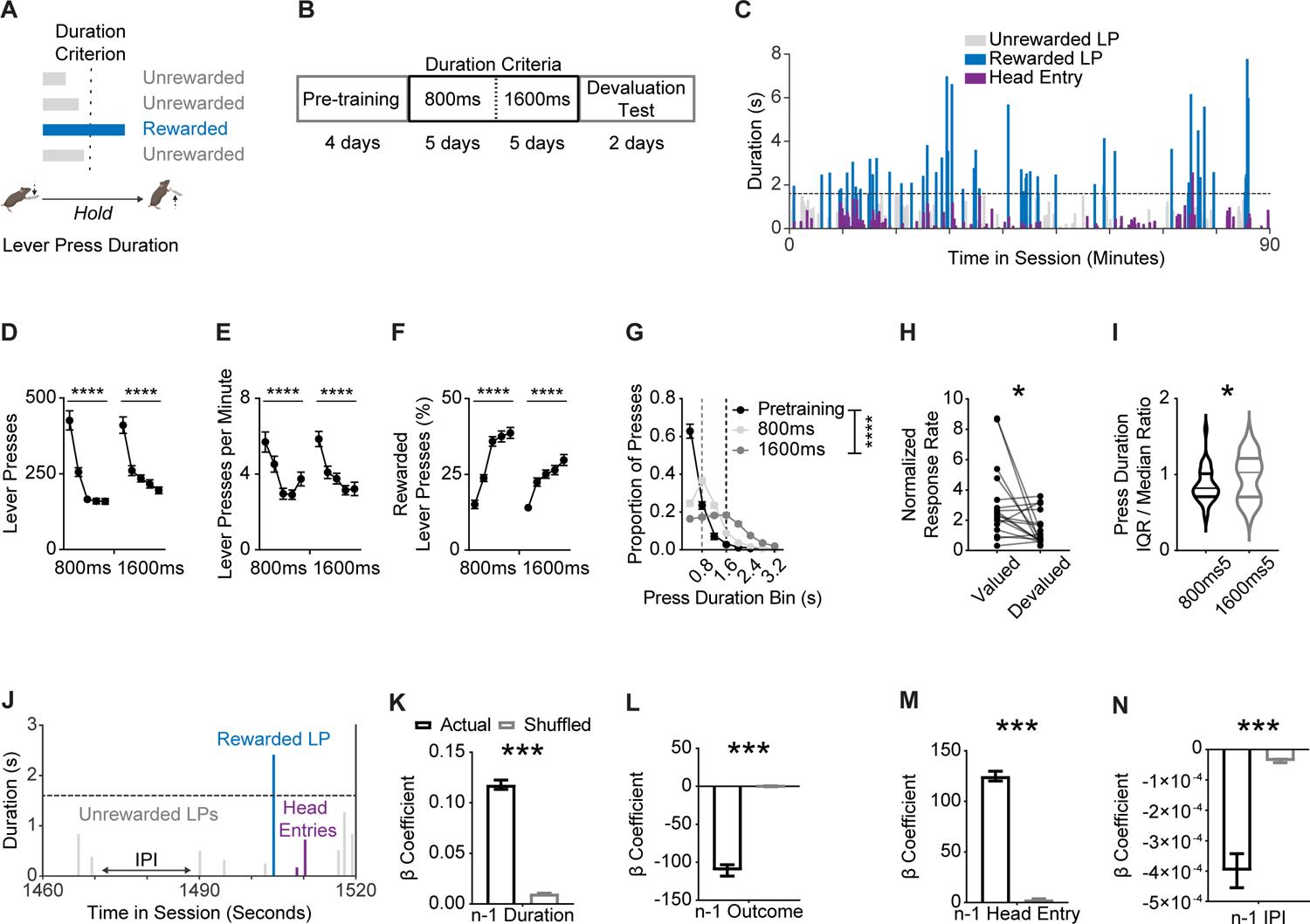
Mice learned to adjust self-paced, self-generated lever pressing actions across inferred contingency and outcome value changes. (A) Behavior schematic demonstrating how mice must press and hold down a lever beyond a minimum duration to earn a food reward. (B) Training schedule for the lever press hold down task. Pretraining sessions were followed by sessions with a minimum duration criterion. Devaluation testing procedures occurred thereafter. (C) Representative data from one mouse showing variability of lever pressing and head entry behavior within a session. Dashed line indicates 1600 ms criterion. (D–E) Total lever presses (D), (E) lever pressing rate and (F) percentage of lever presses that exceeded the duration criterion across sessions. (G) Histogram of lever press durations (400 ms bins) averaged for all pretraining, 800 ms, and 1600 ms duration criterion sessions. (H) Normalized response rates (lever presses per minute) in valued and devalued states throughout devaluation testing. (I) Ratio of lever press duration Interquartile Range (IQR) and median during final 800 ms and 1600 ms duration criterion sessions. (J) Zoomed-in behavior from representative data shown in (C). (K–N) β coefficients of LME model relating current lever press duration (n) to prior (n - 1) press durations (K), press outcome (i.e. was lever press rewarded) (L), head entry (M), and interpress interval (IPI) (N) for actual and order shuffled data. 800 ms and 1600 ms refer to days where the criterion was >800 ms or >1600 ms. Data points represent mean ± SEM. Significance markers in k-n indicate comparisons to order shuffled data. Shuffled data are mean ± SEM of 1000 order shuffled β coefficients. *p < 0.05, ***p < 0.001, ****p < 0.0001, See also Figure S1 and Table S1 for more data.

After initial lever press pre-training, lever press duration criterion was set at >800 ms for five daily sessions, followed by five daily sessions with a >1600 ms duration criterion (Figure 1B). A representative session from a well-trained mouse during a 1600 ms criteria day shows variability in the duration and frequency of lever presses made across the session (Figure 1C). Examining the macroscopic aspects of lever press behavior showed mice reduced the number of total lever presses made (Figure 1D, one-way RM ANOVAs for 800 ms and 1600 ms training durations, Fs > 35.84, ps < 0.0001) and decreased response rates across each duration criteria (Figure 1E, one-way RM ANOVAs Fs > 18.16, ps < 0.0001). Mice increased successful performance in the task within each duration criteria rule, as shown by an increase in the percentage of total presses made that exceeded the minimum duration criterion (Figure 1F; one-way RM ANOVAs for 800 ms and 1600 ms training durations Fs > 31.32, ps < 0.0001). In addition, we observed rightward shifts in the distributions of press durations made when duration criteria shifted, from short durations made in pre training sessions (i.e. no duration requirement to earn a pellet reward) to longer durations made across the 800 ms and 1600 ms duration criterion sessions (Figure 1G; two-way RM ANOVA, main effect of Duration Bin F_1.943, 69.96_ = 336.5, p < 0.0001, main effect of Criterion F_1.067,38.40_ = 7.061, p < 0.05, and an interaction (Duration Bin * Criterion) F_2.622, 94.38_ = 104.8, p < 0.0001). Initiation of lever-press behavior in this task was goal-directed as assayed by outcome devaluation testing procedures (Figure S1A). To account for variance in levels of responding, we normalized response rates during testing to the last two days of training. We observed a reduction in normalized response rates in the devalued compared to valued states (Figure 1H; Paired t-test, t_16_ = 2.482, p < 0.05) (see also Figure S1B). Of note, the percentage of successful lever presses did not differ between valuation states (Figure S1C). This result aligned with prior findings that raise questions about what behavioral mechanisms control remaining lever presses in goal-directed responding.^33, 42^ Thus, broad behavioral performance measures suggested that mice used inferred contingency and expected outcome information to guide their lever press behavior.

However, it was unclear what information mice were using to adjust lever press performance. As previously reported,^33^ the behavior of mice violated the scalar property of timing (ratio of median and interquartile range (IQR) of lever press durations) (Figure 1I; Paired t-test, t_36_ = 2.588, p < 0.05). This suggested that mice may not have exclusively timed each lever press independently, but rather that lever press durations were also influenced by preceding lever press durations as well as other sources of information derived from recent experience, including prior reward delivery, prior checking behavior, as well as the time passed between lever presses (interpress-interval) (Figure 1J). To investigate whether these sources of observable and inferred information influenced lever pressing behavior, we built linear mixed effect models (LMEs) that measured the predictive relationship of these behavioral events on the subsequent lever press duration (n). LME regression coefficients (β) from behavioral covariates of interest were then compared against lever press order-shuffled data via permutation testing.

We found that mice relied on prior experiential information to guide lever pressing. First, sequential lever press durations (n - 1) were related to one another (Figure 1K; p < 0.01, Table S1, top), with the predictive relationship decaying up to the 10th prior (n - 10) lever press duration (Figure S1D; ps < 0.05, Table S1, middle), confirming mice inferred prior lever press durations to adjust future responding. Mice made shorter presses after reward delivery (n - 1 Outcome), potentially indicative of titrating lever press durations for performance success (Figure 1L; p < 0.01).^33, 72^ Checking behavior, indexed via a head entry into the food receptacle, increased the subsequent lever press duration (Figure 1M; p < 0.01). Furthermore, the longer the interval in between presses, the shorter the subsequent lever press duration (Figure 1N; p < 0.01). Importantly, and in line with a prior report,^33^ we found that the relationship between sequential presses (β coefficient for n and n-1) was modified by whether the animal made a head entry (i.e. checking behavior) as well as how much time had elapsed (i.e. interpress interval) between presses (Table S1, bottom). In contrast, reward delivery did not alter the relationship between current (n) and prior (n - 1) lever press durations (Table S1, bottom), suggesting that the presence or absence of reward did not change how mice used prior lever press duration information to guide subsequent performance. The use of this experiential information improved performance. We tested LME model performance using individual session data and found a positive relationship between model R^2^ for an individual and that individual’s overall session performance efficiency (Figures S1E and S1F). Subjects used experiential information to adjust behavior even during early learning; for example mice relied on prior duration information to a similar degree between early (i.e. first few 800ms days) and late acquisition (i.e. last few 1600 ms days) (Figure S1G). However, the influence of other sources of experiential information, such as prior outcome, checking behavior, and interpress interval, increased as training continued (Figures S1H–S1J). The above findings replicate previous results showing that when behavior is largely uninstructed, mice rely on numerous sources of experiential information to guide volitional action control,^33^ including inferences about prior action performance.

### lOFC populations differentially encode actions and action-related information

We next sought to investigate whether OFC may reflect the use of such experiential information during decision-making, particularly lever press duration information. As head fixation can have large effects on context-dependent behaviors,^76^ we monitored OFC population Ca^2+^ activity of CaMKII+ projection populations (rAAV5/PAAV-CaMKIIa-GCaMP6s) using *in vivo* fiber photometry in freely-moving mice (C57BL/6J, n = 9 mice, 6 males, 3 females) as they performed the lever-press hold down task during 1600 ms duration criterion training (Figures 2A and S2). A perievent histogram of Ca^2+^traces ordered by press duration revealed that CaMKII+ OFC projection population activity was modulated at select epochs relative to lever press initiation and execution (Figure 2B). We segmented CaMKII+ fluorescence activity traces by whether or not the lever press was eventually rewarded. Group averaged CaMKII+ OFC projection population activity was modulated prior to the onset of a lever press, similarly to previously reported single-unit recordings.^40, 42^ Permutation testing ^77^ revealed that pre onset lOFC Ca^2+^ activity differed with respect to its eventual outcome (Figure 2C; ps < 0.05). Future success-related differences persisted during the lever press (Figure 2D; ps < 0.05) and success-related differences were observed after the lever press release (Figure 2E; ps < 0.05). Indeed, we observed greater levels of CaMKII+ OFC projection population activity in head entries that followed a successful lever press than in head entries following an unsuccessful lever press, corresponding to a time point during which mice had access to food pellet-related sensory and consummatory information (Figure 2F; ps < 0.05).

**Figure 2.**
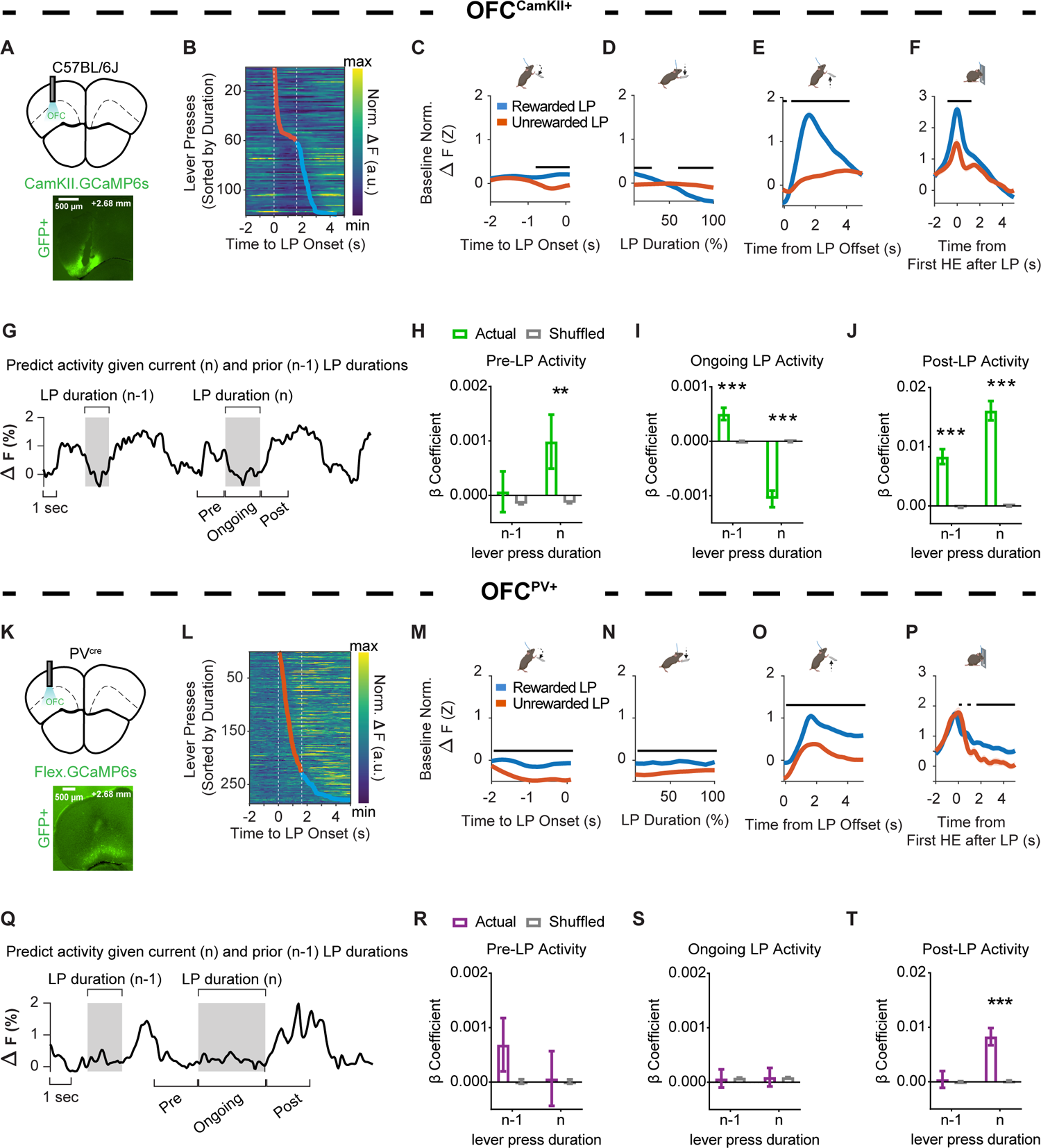
OFC^CamKII+^, but not OFC^PV+^, Ca^2+^ activity encodes prior action information. (A) (top) Anatomical schematic and (bottom) representative histology of (A) OFC^CamKII+^ *in vivo* Ca^2+^ fiber photometry experiments. Approximate bregma coordinate AP +2.68 mm. (B) Representative data heat map of OFC^CamKII+^ normalized fluorescence changes relative to lever press initiation, ordered by lever press duration. (C–F) Ca^2+^ activity from OFC^CamKII+^ populations z-scored normalized relative to a pre-lever press onset baseline period and aligned to (C) lever press onset, (D) lever press duration (presented as the relative percentage of total lever press duration), the (E) offset (i.e. termination) of a lever press, and the (F) first head entry made after a lever press. (G) Representative traces indicating changes in OFC^CamKII+^ Ca^2+^ fluorescence over time. (H–J) β coefficients from LME models relating OFC^CamKII+^ Ca^2+^ activity to current and prior durations for actual and order shuffled data (H) before press onset (Pre-LP Activity), (I) during the press (Ongoing LP Activity), and (J) after press offset (Post-LP Activity). (K) (top) Anatomical schematic and (bottom) representative histology of OFC^PV+^ in vivo Ca^2+^ fiber photometry experiments. Approximate bregma coordinate AP +2.68 mm. (L) Representative data heat map of OFC^PV+^ normalized fluorescence changes relative to lever press initiation, ordered by lever press duration. (M–P) Ca^2+^ activity from OFC^PV+^ populations z-scored normalized relative to a pre-lever press onset baseline period and aligned to (M) lever press onset, (N) lever press duration (presented as the relative percentage of total lever press duration), the (O) offset (i.e. termination) of a lever press, and the (P) first head entry made after a lever press. (Q) Representative traces indicating changes in OFC^PV+^ Ca^2+^ fluorescence over time. (R–T) β coefficients from LME models relating OFC^PV+^ Ca^2+^ activity to current and prior durations for actual and order shuffled data (R) before press onset (Pre-LP Activity), (S) during the press (Ongoing LP Activity), and (T) after press offset (Post-LP Activity). LP = lever press, HE = Head Entry. Black lines in C–F and M–P indicate significant differences between Rewarded and Unrewarded lever presses via permutation testing (p < 0.05). For B and L, dashed white lines indicate 1600 ms session criterion window. Orange markers indicate the termination of an unrewarded lever press. Blue markers indicate the termination of rewarded lever press. For G and Q, shaded regions indicate the duration of the current (n) or prior (n - 1) lever press. Ca^2+^ activity magnitude predicted by LME models includes −2s to 0s before (Pre), during (Ongoing), or 0 to 2s after (Post) the current lever press. Shuffled data are the mean ± SEM of 1000 order shuffled β coefficients. ** p < 0.01, *** p < 0.001. See also Figure S2, Table S2 and Table S3 for more data.

Our results suggest that excitatory projection neurons in OFC can reflect information related to eventual performance outcomes before and throughout an instrumental action. To investigate whether this information originated from aspects of experiential information, we built LME models which aimed to predict lever press aligned changes in calcium activity given the current lever press duration and prior experiential information, with a focus on action-related information (Figure 2G). Prior to lever press onset, we found a significant relationship between CaMKII+ OFC projection population activity and the upcoming (n) lever press duration (Figure 2H). This significant relationship was also found while the animals held down the lever (Figure 2I) and was still present at termination of the lever press (Figure 2J). In other words, prior to lever press onset, greater increases in lOFC CaMKII+ activity were associated with longer durations of the upcoming action. However, during lever press execution, lower levels of activity corresponded to longer lever presses. At lever press offset, longer lever presses were associated with increased lOFC CamKII activity. In contrast, when we examined whether lOFC CamKII+ activity reflected prior (n - 1) action-related information (i.e. prior press duration), we found no relationship between the prior lever press duration and current lOFC CamKII+ calcium activity at lever press onset (Figure 2H), suggesting lOFC CamKII+ neurons are not maintaining prior duration information. However, there were significant relationships between current lOFC CamKII+ calcium activity and prior (n - 1) lever press during lever press execution (Figure 2I) as well as at lever press offset (Figure 2J). We also found that CaMKII+ OFC projection population activity during these lever press epochs was modulated by whether the prior lever press was rewarded or not, whether a checking head entry was made, as well as the time from prior lever press (Table S2). Together, our results suggest that current and prior action-related information as measured by lever press durations, as well as broader experiential information, is differentially encoded by CamKII+ projection populations in the OFC. This encoding occurred during action initiation and execution and further suggests that OFC projection circuits may be recruited for action information during adaptive behavior.

OFC projection circuits do not act in isolation. Local GABAergic interneurons are crucial for the control of local circuit inhibition^68, 70, 71^ with PV+ interneurons playing a critical role in tuning spike timing and synchronizing network oscillations.^67, 69^ As cortical projection and local inhibitory populations receive long-range cortical input,^64, 65^ it may be that PV+ lOFC inhibitory population activity reflects experiential information similar to that of CaMKII+ lOFC projection populations. However, it could also be that PV+ lOFC inhibitory population recruitment supports computations performed by local OFC projection populations with less regard to any specific type of information. Therefore we performed fiber photometry experiments monitoring Ca^2+^ activity of virally targeted PV+ lOFC interneuron populations (rAAV5/pAAV.CAG.Flex.GCaMP6s.WPRE.SV40) in freely moving PV^cre^ mice as they performed the lever-press hold down task (PV^cre^, n = 8, 5 males, 3 females) (Figure 2K). A peri-event histogram of baseline normalized PV+ traces ordered by press duration suggested that PV+ lOFC interneuron population activity was modulated at select epochs relative to lever press initiation and execution, but with patterns that notably differed from CaMKII+ lOFC projection populations (Figure 2L). Permutation testing revealed that group averaged lOFC PV+ Ca^2+^ activity was modulated prior to the onset of a lever press, with smaller reductions of activity observed with lever presses that would be rewarded (Figure 2M). Reward-related differences in lOFC PV+ Ca^2+^ activity persisted during ongoing lever press execution (Figure 2N) and after lever press offset (Figure 2O), increasing to a larger degree for rewarded than unrewarded press durations. In contrast to CaMKII+ lOFC projection populations, lOFC PV+ Ca^2+^ activity associated with head entries made following lever press release reached similar levels regardless of the presence of a reward (Figure 2P).

To investigate whether PV+ lOFC inhibitory population activity is modulated by aspects of experiential information, we again built LME models which aimed to predict lever press aligned changes in PV+ Ca^2+^ activity given prior and current lever press durations, as well as other sources of experiential information (Figure 2Q). In contrast to the CaMKII+ lOFC projection population, PV+ lOFC interneuron population Ca^2+^ activity prior to (Figure 2R) and throughout the lever press (Figure 2S) was not predictive of ongoing (n) or prior (n - 1) lever press durations, prior checking behavior, nor the inter-press interval, but was predictive of whether the prior press was rewarded or not (Table S3).

However, at lever press offset, PV+ lOFC interneuron population activity was predictive of the duration of the lever press that was just executed (n), as well as prior checking behavior and lever press outcome (Figure 2T and Table S3). Our results suggest that, unlike CamKII+ projection populations, PV+ inhibitory population activity in lOFC largely reflects outcome-related information during lever pressing and consequence-related information after lever press termination. Together, our results suggest that inferred action information is differentially encoded by lOFC sub-populations.

### lOFC uses action-related information to modify behavior

That lOFC populations can reflect prior and current lever press durations suggests that lOFC activity may functionally contribute to the use of action information to adjust behavior. To test this hypothesis, we first aimed to selectively disrupt local OFC activity in a temporally-specific manner during lever press execution. We used an optogenetic approach to bilaterally activate PV+ lOFC inhibitory populations with an excitatory opsin (rAAV5/Ef1a-DIO-hChR2(H134R)-eYFP) to inhibit local lOFC projection population activity (PV^cre^; n = 6 ChR2, 5 males, 1 female; n = 8 YFP, 5 males, 3 females) (Figure 3A).^29, 78^ Optical stimulation of PV+ OFC inhibitory populations was behaviorally-dependent on the execution of a lever press, such that the initiation of every lever press, independent of their eventual duration, triggered light delivery (470 nm 20 Hz, 5 ms pulses) that continued until press termination (Figure 3B). Stimulation days occurred after task acquisition (Figure 3C). In days in which light was delivered, ChR2 mice maintained similar rates of responding (Figure 3D; p > 0.05), but reduced the percentage of rewarded lever presses compared to fluorophore controls (Figure 3E; two-way RM ANOVA, main effect of Treatment only F_1, 12_ = 4.990, p < 0.05). Comparisons of lever press duration distributions suggested that light activation altered the distribution pattern of lever press durations in ChR2 mice (Figure 3F; two-way RM ANOVA, main effect of Duration Bin F_1.813, 21.76_ = 56.73, p = 0.0001; marginally significant interaction (Duration Bin * Treatment) F_9, 108_ = 1.967, p = 0.05). The above data suggest disruption of lOFC activity during action execution impaired successful performance.

**Figure 3.**
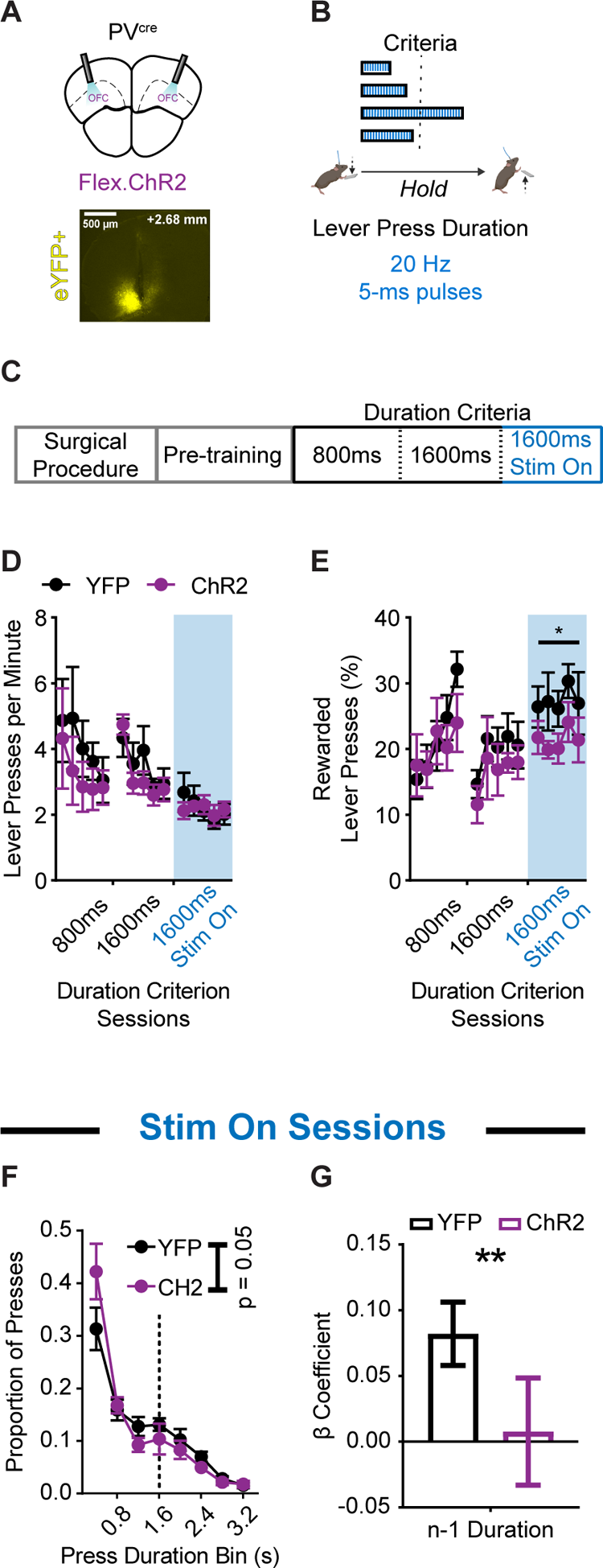
Optogenetic excitation of OFC^PV+^ populations during action execution reduces rewarded performance and use of prior action information. (A) (top) Schematic and (bottom) example histology of ChR2 optogenetic excitation of OFC^PV+^ neurons. Approximate bregma coordinate AP +2.68 mm. (B) Behavior schematic demonstrating how 470 nm light delivery (20 Hz, 5 ms pulses) occurred for the duration of every lever press made. (C) Training schedule for optogenetic experiments. Pretraining sessions were followed by sessions with a minimum duration criterion. Sessions in which light was delivered occurred thereafter. (D and E) Lever pressing rate (D) and (E) percentage of lever presses that exceeded the duration criterion across sessions. Blue shaded region indicates the sessions in which light was delivered. (F) Histogram of lever press durations (400 ms bins) averaged for all 1600 ms duration criterion sessions during which light was delivered. (G) β coefficients of LME model relating current lever press duration (n) to prior (n - 1) press durations for YFP and ChR2 cohort actual data. Significance marker indicates comparisons to 1000 group shuffled data. 800 ms and 1600 ms refer to days where the criterion was >800 ms or >1600 ms. 1600 ms Stim On refers to days where criterion was >1600 ms and light was delivered. Data points represent mean ± SEM. *p < 0.05, **p < 0.01. See also Table S4 for more data.

We next examined whether this disruption of OFC activity impaired performance in part by changing the predictive relationship between prior (n - 1) and ongoing (n) lever press durations. We added a term to our behavioral LME model that accounted for the presence or absence of the excitatory opsin (Treatment) in each animal. We found a significant interaction between the prior lever press duration and the presence of the opsin (Duration_n-1_ * Treatment) in predicting subsequent lever press durations (Table S4; p < 0.05). A representation of group-segmented β coefficients showed a reduced relationship between prior (n - 1) and current (n) lever press durations in ChR2 animals compared to fluorophore controls (significant compared to 1000 group-shuffled data) (Figure 3G; p < 0.01). The above suggests lOFC activity supports action-related information.

However, disrupting lOFC activity during every lever press as described above prevented us from determining whether lOFC activity contributed to the encoding of action information, or to the use of prior action information to guide ongoing performance. Another possibility is that inhibiting lOFC during the lever press was akin to enhancing the general reduction in activity normally observed when the animal holds down the lever (Figure 2D),^42^ thereby facilitating lOFC processes that could be competing with other action-related processes and associated circuits. Furthermore, inducing interneuron mediated inhibition during every lever press may have recruited compensatory mechanisms for task performance, such that the observed behavioral effects may not be directly attributable to a loss of lOFC function. Therefore, we next directly examined the acute behavioral contributions of lOFC activity patterns and their support for the encoding and/or use of action-related information. We bilaterally expressed an excitatory opsin in CamKII+ projection neurons (rAAV5/CamKII-hChR2(H134R)-eYFP-WPRE) to induce non-physiological increases in lOFC activity during lever pressing (i.e., when activity is normally decreased) in a small subset of actions made during the session. In mice (C57BL/6J; n = 7 ChR2, 5 males, 2 females; n = 5 YFP, 2 males, 3 females) (Figure 4A), we paired light activation (470 nm 20 Hz, 5 ms pulses) only to every 7th lever press. This allowed us to examine whether proper patterning of lOFC activity was necessary for the encoding (i.e. would light activation during prior (n - 1) press would prevent its encoding and therefore reduce its contribution to the ongoing (n) duration?) and/or subsequent use of action-related information (i.e. would light activation during the ongoing (n) press prevent the use of prior (n - 1) duration information?). Light stimulation continued up until the lever press was terminated (Figure 4B) and stimulation occurred post task-acquisition (Figure 4C).

**Figure 4.**
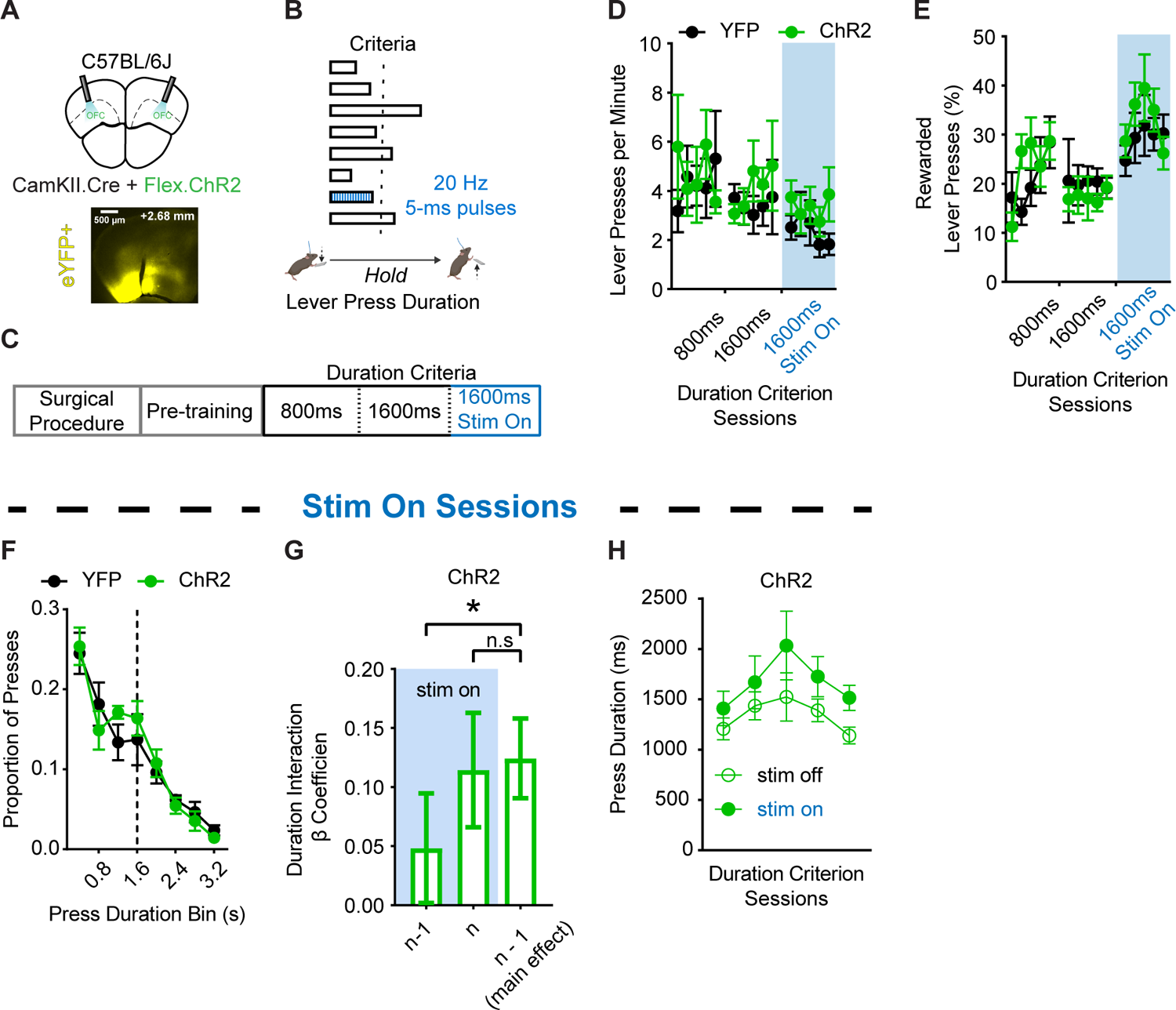
Selective optogenetic excitation of OFC^CamKII+^ populations during action execution does not impair performance but affects use of prior action information. (A) (top) Schematic and (bottom) example histology of ChR2 optogenetic excitation of OFC^CamKII+^ neurons. Approximate bregma coordinate AP +2.68 mm. (B) Behavior schematic demonstrating how 470 nm light delivery (20 Hz, 5 ms pulses) occurred for the duration of every 7th lever press made. (C) Training schedule for optogenetic experiments. Pretraining sessions were followed by sessions with a minimum duration criterion. Sessions in which light was delivered occurred thereafter. (D and E) Lever pressing rate (D) and (E) percentage of lever presses that exceeded the duration criterion across sessions. Blue shaded region indicates the sessions in which light was delivered. (F) Histogram of lever press durations (400 ms bins) averaged for all 1600 ms duration criterion sessions during which light was delivered. (G) β coefficients of post-hoc LME model relating current lever press duration (n) to prior (n - 1) or current (n) press durations for ChR2 cohort actual data. Significance marker indicates comparisons to 1000 order shuffled data. (H) Lever press durations of 1600 ms duration criterion sessions during which light was delivered on every 7th lever press, segmented by whether presses were paired with light activation or not. 800 ms and 1600 ms refer to days where the criterion was >800 ms or >1600 ms. 1600 ms Stim On refers to days where criterion was >1600 ms and light was delivered. Data points represent mean ± SEM. *p < 0.05, *** p < 0.001. See also Figure S3 and Table S5 for more data.

Both ChR2 and YFP mice reached similar rates of lever pressing and performance throughout training, including the 5 daily sessions during which light was delivered (Figures 4D and 4F, ps > 0.05), suggesting that acute disruptions of lOFC activity during action execution on a small subset of lever presses did not affect gross performance measures. A behavioral LME model that accounted for the presence or absence of the excitatory opsin in each animal found a significant interaction between the presence of the opsin and the predictive relationship between current and prior lever press durations (Duration_n-1_ * Treatment) (Table S5, top, p < 0.001). To determine the direct effects of stimulation on the predictive relationship between prior (n - 1) and ongoing (n) lever press durations, we built post-hoc LME models using either the ChR2 or YFP datasets that accounted for the presence or absence of light stimulation in each lever press. We found a significant interaction between prior press stimulation and the predictive relationship between current and prior press durations (Duration_n-1_ * Stimulation_n-1_) in ChR2-expressing mice that was absent in fluorophore control mice (Table S5, middle and bottom). Inspection of the LME model interaction β coefficients in ChR2 mice data showed light activation during the n - 1 lever press reduced the contribution of n-1 lever press duration from informing the ongoing (n) lever press (p < 0.05) (Figure 4G; left and right bar comparison). However, light activation during the ongoing (n) lever press did not reduce that lever press’s reliance on n - 1 duration information (p > 0.05) (Figure 4G; middle and right bar comparison). These data suggest the proper patterning of lOFC activity supports processes related to the encoding (i.e. significant effect on n - 1 press activation), but not the retrieval and use (i.e. non-significant effect on n press activation) of action information to guide future action execution. Light activation did not induce selective decreases or increases to the lever press duration itself (Figure 4H; p > 0.05). In addition, light activation did not change the proportion of successful lever presses (Figure S3A) nor the time it took to initiate the subsequent lever press (Figure S3B). In conjunction with our PV+ inhibitory population disruptions, these data suggest proper patterning of lOFC activity supports processes related to the encoding of action information to guide future action execution.

### Loss of functional lOFC circuit increases reliance on immediate prior actions and outcomes

Different action strategies can be used to achieve the same goal.^1, 3, 79, 80^ When one circuit is offline another may be recruited to support decision-making and adaptive behavior (i.e the emitted behavior does not reflect the function of the perturbed circuit).^40, 57–59^ Within this framework, chronic removal of OFC circuits, as is often done in lesion studies, could bias recruitment of compensatory or parallel mechanisms for volitional action control. To test how chronic lesions to lOFC projection neurons would impact lever-press hold down task performance, prior to training we bilaterally ablated OFC CamKII+ neurons using a cre-dependent caspase approach that committed infected neurons to apoptosis (rAAV5/AAV-Flex-taCasP3-TEVP) (C57BL/6J; Lesion n = 16, 13 males, 3 females; Sham n = 23, 14 males, 9 females) (Figures 5A, 5B and S4A).^81, 82^ LOFC lesions improved task efficiency. lOFC lesioned mice had lower overall response rates than Sham mice during 1600 ms training (Figure 5C; two-way RM ANOVA, main effect of Session F_1.912, 70.73_ = 16.34, p < 0.0001 and an interaction (Session * Treatment) F_4, 148_ = 2.875, p < 0.05). However, lesioned mice showed a higher percentage of rewarded lever presses than Sham mice (Figure 5D; two-way RM ANOVA, main effect of Session F_2.347, 86.83_ = 22.55, p < 0.0001, and Treatment F_1, 37_ = 6.804, p < 0.05). Furthermore, lOFC-lesioned mice showed a rightward shift in the distribution of lever press durations (Figure 5E; two-way RM ANOVA, main effect of Duration Bin F_2.872, 106.2_ = 98.52, p < 0.0001 and an interaction (Duration Bin * Treatment) F_9, 333_ = 2.336, p < 0.05) throughout the 1600 ms duration criterion sessions. A behavioral LME model that accounted for the presence or absence of the lesion in each animal found a significant interaction with treatment group reflecting an altered relationship between the current (n) and prior (n - 1) lever press durations (Duration_n-1_ * Treatment) (Table S6; p < 0.001). Treatment group-segmented β coefficients showed a larger positive relationship between prior and subsequent durations in Lesion animals compared to Sham controls (significant compared to 1000 group-shuffled data) (Figure 5F, p < 0.001). We also found a significant interaction between the outcome of the prior lever press and lesion group (Outcome_n-1_ * Treatment) (Table S6; p = 0.001). A representation of treatment group-segmented β coefficients showed a greater negative relationship between prior outcome and subsequent durations in Lesion mice compared to Sham mice (significant when compared to 1000 group-shuffled data) (Figure 5G; p < 0.001). The effect of lesions on use of prior action and outcome information was strongest during early 800 ms and early 1600 ms training (Figures S4B–S4E), suggesting that OFC lesions led to recruitment of other circuits which relied on immediate action and outcome history to a greater extent when contingencies were increased.

**Figure 5.**
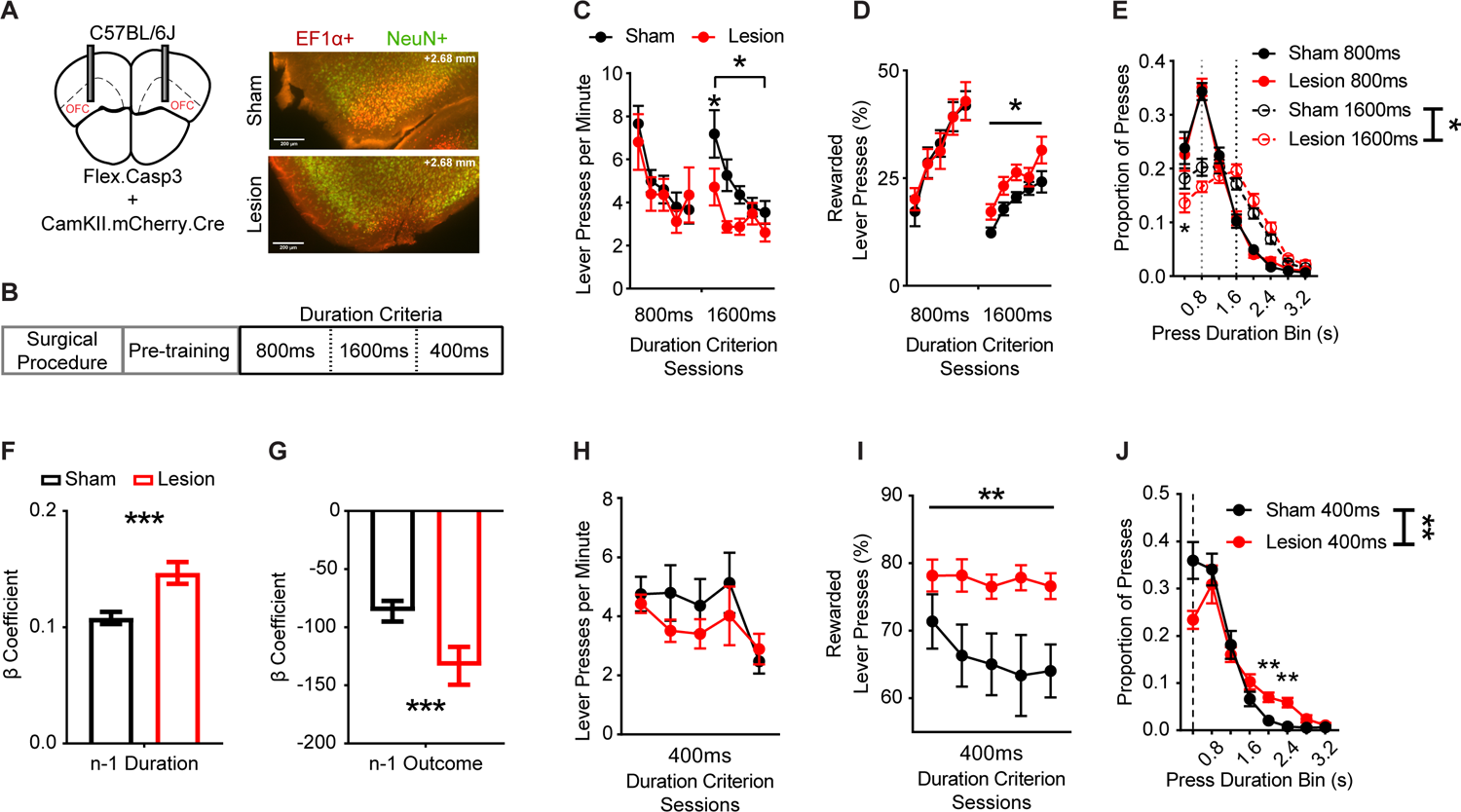
Pretraining OFC^CamKII+^ lesions increase rewarded performance and use of prior action information. (A) (left) Schematic and (right) example histology of sham and Cre-dependent caspase lesion of OFC neurons. Red indicates AAV-EF1α-DIO-mCherry expression. Green indicates immunohistochemical reactions for neural nuclear protein NeuN. Approximate bregma coordinate AP +2.68 mm. (B) Training schedule for lesion experiments. Pretraining sessions were followed by sessions with a minimum duration criterion. (C and D) Lever pressing rate (C) and (D) percentage of lever presses that exceeded the duration criterion across sessions. (E) Histogram of lever press durations (400 ms bins) averaged for all 800 ms and 1600 ms duration criterion sessions. Dotted lines indicate 800 ms (grey) and 1600 ms (black) duration criteria. (F and G) β coefficients of LME model relating current lever press duration (n) to prior (n - 1) press durations (F) and press outcome (i.e. was lever press rewarded) (G) for Sham and Lesion cohort actual data. Significance markers indicate comparisons to 1000 group shuffled data. (H and I) Lever pressing rate (H) and (I) percentage of lever presses that exceeded the 400 ms duration criterion across sessions. (J) Histogram of lever press durations (400 ms bins) averaged for all 400ms duration criterion sessions. Dotted line indicates 400 ms duration criteria. 400 ms, 800 ms,1600 ms refer to days where the criterion was >400 ms, >800 ms or >1600 ms. Data points represent mean ± SEM. *p <0.05, ** p < 0.01, *** p < 0.001. See also Figure S4 and Table S6 for more data.

lOFC lesions have been reported to reduce behavioral flexibility and impair sensitivity to rule reversals.^17, 23, 46, 83–85^ We conducted an additional 5 daily sessions in which the duration criterion was reduced to 400 ms for a subset of animals (C57BL/6J; Lesion n = 16, 13 males, 3 females; Sham n = 8, 6 males, 2 females). We found that while Sham and Lesion mice showed similar rates of lever pressing (Figure 5H; two-way RM ANOVA, main effect of Session only F_1.645, 36.20_ = 4.274, p < 0.05), Lesion mice performed more efficiently than Sham mice as indexed by a higher percentage of rewarded lever presses (Figure 5I; two-way RM ANOVA, main effect of Treatment only F_1, 22_ = 1.590, p < 0.01) and showed a rightward shift in the distribution of lever press durations (Figure 5J; two-way RM ANOVA, main effect of Duration Bin F_2.369, 52.13_ = 70.97, p < 0.0001 and an interaction (Duration Bin * Treatment) F_9, 198_ = 3.164, p < 0.01). The above suggests mice with lOFC lesions did not adjust their performance to the same degree as sham animals when the duration contingency was reduced in duration. Instead, lOFC lesion mice continued to perform longer lever presses, a strategy that improved efficiency but differed from the exploration of effort that intact mice exhibited.

## Discussion

Here we identify a novel role for lOFC in action control. By examining adjustments to volitional actions within the continuous context in which they occur, we were able to separate control processes dictated by inferences about prior actions from those dictated by inferences of action-outcome contingency or expected outcome value. In doing so, we saw clear evidence that mice recruit and use lOFC activity to encode action-related information that can be used for inferences critical to adaptive control of behavior. A loss of lOFC circuits left mice more reliant on a strategy of repeating action execution to gain reward and impaired action exploration and the updating of action contingencies. This raises the hypothesis that lOFC circuit disruptions seen in psychiatric disorders may give way to compensatory mechanisms that promote repetitive action control by exploiting the reliance on learned rules, even when disadvantageous.

There is increasing evidence of lOFC disruption in psychiatric disorders characterized by disrupted action control, including substance use disorders and compulsive disorders,^86–89^ highlighting the need for a greater understanding of OFC’s contribution to the use of action-related information. Actions made during decision-making are often autonomous and unconstrained, occurring in contexts in which contingencies and associative structure of ongoing tasks are partially observable at best.^3, 4, 36, 90^ We show that eschewing trial-based, cued choice structure in task design has advantages for understanding how adaptive behavior is realized when actions are self-paced, self-generated, and mainly reliant on inferences that recruit information from past experiences (Figure 1). The unstructured, self-paced nature of our task allowed us to identify patterns of lOFC activity that reflected prior action information as well as other sources of experience. As OFC populations have been shown to process cue-related and reward-predictive information,^16, 20–22, 25, 26, 91^ our findings suggest that OFC can also represent and encode action-related information to guide adaptive behavior depending on the behavioral context.

While prior single unit recordings from largely unclassified populations have shown lOFC neurons can reflect sensory, predictive, and outcome-related information,^20–22, 48^ here we find that lOFC populations appear to be differentially recruited to support encoding of action-related information. CamKII+ projection neuron activity increased prior to the onset of the action, decreased during action execution, and ramped back up after the action was terminated (Figure 2). These lOFC activity patterns were reminiscent of prior single-unit recording activity observations in rodents performing the same task,^42^ suggesting single neuron population activity likely tracks population calcium activity. Furthermore, lOFC excitatory projection neuron activity reflected current and prior action-related information during ongoing action execution. Temporally precise and behavioral dependent perturbation to these endogenous lOFC CamKII+ neuron activity patterns decreased reliance on action-related information. Intriguingly, these lOFC CamKII+ activity patterns differed in timing and magnitude compared to PV+ interneuron activity patterns. PV+ populations showed relatively little performance-dependent modulation prior to and throughout action execution, little outcome encoding during reward checking behaviors, and maintained little representation of action-related information. While the use of population calcium measurements may not capture individual neuron encoding of action-related information, these findings do suggest that action-related information is reflected in recruitment of lOFC CamKII+ populations and to a much lesser extent PV+ populations. Cortical PV+interneurons are thought to gate information flow within cortical microcircuits through spike-timing enforcement of projection neuron firing.^68–71^ Perhaps the observed differential patterns of lOFC activity are reflective of local microcircuit interactions that facilitate the flow of action-related information through this region.

OFC has been hypothesized to integrate and relay information from prior experiences to its broader circuits to support ongoing decision-making processes.^6, 10^ lOFC neurons have been shown to encode prior reward-predictive information before subsequent choices are made.^20–22^ Here we observed lOFC CamKII+ activity reflected the future success or failure of the imminent and ongoing lever-press (Figure 2). Furthermore, LME modeling showed that this activity reflected and maintained information related to the prior lever press duration (Figure 2). However, optogenetic perturbation to lOFC populations during ongoing action execution revealed that this lOFC activity encodes action information but such activity is not important for using action history to guide current performance (Figures 3 and 4). Such types of information within OFC populations has been hypothesized to be influenced by broader circuit innervation, such as the ventral tegmental area^16^ and the basolateral amygdala.^91^ Investigations examining what OFC relays to downstream targets suggests that lOFC terminals in the dorsal striatum convey both action and outcome-related information to influence ongoing behaviors.^45, 92–94, 95^ In addition, lOFC projections convey rule information to premotor cortical areas used to guide arbitration between rule exploration and exploitation.^33, 96, 97^ However, not all prefrontal cortical areas are likely to play the same role for action control. Indeed, we have shown that unlike lOFC, the premotor cortex and its output to the dorsal striatum is involved in the use of action history to guide performance.^33^ As task-related information has been seen broadly across the cortex,^98^ how specific experiences recruit different cortical and subcortical activity patterns should be explored in the future.

The loss of lOFC CamKII+ projection neuron populations did not lead to a loss of efficacy in performance or action control. Instead, lesioned mice showed more efficacious performance and persisted in making longer lever presses despite the change to a shorter duration criterion for success (Figure 5). Historically, OFC lesions have been thought to spare behavior that relies on observable information while impairing behavior when task demands get more abstract and begin to rely on inference-based information.^5^ Our results suggest a nuanced view of what lOFC may contribute to action control. While lOFC may not be necessary for direct action control per se, it does appear to be recruited when behavioral control recruits the inclusion of action and outcome-related inferences, broad experiences, and exploration.^38^ Perhaps a functional loss of OFC circuits engaged compensatory mechanisms (e.g., recruitment of other circuits) that biased control of behavior to rely on more immediate sources of reward-related information.^79, 80, 99,100,101^ In other words, lesioned animals may not have favored a shift in lever pressing strategy since long durations were still producing rewards. In addition, lesioned mice may have reduced the degree of exploration normally exhibited. Both hypotheses suggest lOFC CamKII+ projection neuron lesions left mice repeating actions to exploit a known rule. Thus activity originating from lOFC appears to support adaptive control of behaviors that in its absence facilitate action strategies that rely on similar behaviors if they had been previously successful.

OFC dysfunction is found in disease states associated with repetitive behaviors and disrupted action control, such as in obsessive compulsive disorder and substance use disorders.^88, 89^ Investigating how information derived from past behaviors are integrated in OFC to influence subsequent actions can aid our understanding of how substances of abuse are sought out and consumed based on prior experience. In humans, repeated transcranial magnetic stimulation studies targeting downstream OFC activity have been shown to be effective at reducing compulsivity.^102, 103^ Here we establish that lOFC population-specific activity can encode action-related information to influence future action implementation. While OFC neurons have been shown to have less action-related recruitment and activity modulation during motor responding compared to some other cortical areas,^104^ discounting its role in processing action information in its entirety limits much needed investigations. The prior experimental discord over whether OFC contributes to action control may have arisen from the use of task parameters that were unable to isolate processes underlying action control from their relationship with associated outcomes or from examining choice behaviors that can use readily observable information. Such tasks may have also elicited a training-induced bias in recruitment of alternative mechanisms for action control. Thus, our findings support the hypothesis that lOFC circuits contribute to adaptive behaviors that rely on prior experience, and that their disruption may lead to an alteration of volitional action control biased towards excessive repetition.

## Acknowledgements

This work was funded by F99-NS120434 (C.C.), F31AA027439 (D.C.S.), R01AA026077 (C.M.G.), and a Whitehall Foundation Award (C.M.G). Some figures included schematics created with BioRender.com.

## Author Contributions

C.C.: Conceptualization, Formal analysis, Funding acquisition, Investigation, Methodology, Visualization, Writing - original draft, Writing - review and edition. D.C.S.: Formal analysis and Writing - review and editing. M.L.V.: Investigation and Writing - review and editing. C.M.G.: Conceptualization, Methodology, Supervision, Funding acquisition, Project administration, Visualization, Writing - original draft, Writing - review and editing.

## STAR Methods

### Resource availability

#### Lead contact

Further information and requests for resources and reagents should be directed to and will be fulfilled by the Lead Contact, Christina M. Gremel, Ph.D.

#### Materials availability

This study did not generate new unique reagents.

#### Data and Code Availability

- The data reported in this paper will be shared by the lead contact upon request.
- All original code has been deposited at GitHub and is publicly available as of the date of publication online in the link listed in the key resources table. All scripts/functions were executed using Matlab 2019a.
- Any additional information required to reanalyze the data reported in this paper is available from the lead contact upon request.

### Experimental model and subject details

C57BL/6J (n = 83, 58 males, 25 females) and PV^cre^ (Pvalb^tm1(cre)Arbr^: n = 36, 26 males, 10 females) mice (>7 weeks/50 PND) (The Jackson Laboratory, Bar Harbour, ME) were housed two to five per cage under a 14:10 hour light:dark in a temperature- and humidity-controlled room and had access to water *ad libitum*. Prior to behavioral procedures, mice were food restricted to 85-90% of their baseline weight for at least 2 days, and were fed a minimum of one hour after daily training (Labdiet 5015). Exploratory analyses for sex and genotype differences in the behavioral cohort revealed similar levels of behavioral performance, and thus data was collapsed across sex and across genotype. Mice were at least 6 weeks of age prior to surgical procedures. All experiments were approved by the University of California San Diego Institutional Animal Care and Use Committee and were carried out in accordance with the National Institutes of Health (NIH) “Principles of Laboratory Care”. Investigators were not blind to the experimental groups. The Animal Care and Use Committee of the University of California, San Diego approved all experiments and experiments were conducted according to the National Institutes of Health (NIH) “Principles of Laboratory Care” guidelines.

## Method Details

### Behavioral Procedures

Daily mouse training sessions occurred within sound attenuating operant chambers (Med-Associates, St Albans, VT) where lever presses (location counterbalanced, either left or right of the food magazine) were required for a reward outcome of regular ‘chow’ pellets (20 mg pellet per reinforcer, Bio-Serv formula F0071). On the first day of pre-training, mice were trained to retrieve pellets from the food magazine (no levers present) on a random time (RT) schedule, with a pellet outcome delivered on average every 120 seconds for 60 minutes. For the next 3 days of pre-training, lever presses were rewarded on a continuous reinforcement (CRF) schedule for up to 15 (CRF15), 30 (CRF30) or 60 (CRF60) pellet reward deliveries or until 90 minutes had passed. For surgical implant experiments, an additional CRF60 training day (for a total of 4 CRF days) was administered with the implant connected to habituate the animal to the tethered connection. Before each session in which the animal was tethered to a fiber optic cable, mice were exposed to a brief (< 60 seconds) bout of low-dose isoflurane anesthesia to connect the ferrule implant. To avoid confounding effects of anesthesia on brain activity, mice were then moved into the procedure room and monitored for a minimum of 30 min before placing them in the operant chamber and initiating the session. The start of each session triggered house-light illumination and the extension of the lever unless stated otherwise.

Following pre-training, mice were introduced to the hold down task. Lever presses now had a duration requirement, such that mice had to continue holding down the lever press for a fixed minimum amount of time in order to earn a pellet reward. Reward delivery occurred only after the termination of a lever press that exceeded the session’s minimum duration criteria, which began with > 800 ms for 5 daily sessions, followed by > 1600 ms for another 5 daily sessions. Each session ended when 90 minutes had elapsed or the mouse had earned 60 total reinforcers, at which point the house light turned off and the lever was retracted. Each lever press onset and termination was timestamped at a 20 ms time resolution to calculate its duration, along with pellet delivery and the start and end of head entries into the food magazine.

### Outcome Devaluation Testing

A subset of the behavioral cohort (n = 18, 10 males, 8 females) was habituated to a novel cage and 20% sucrose solution for 1 hour each day. Following the last day of hold down training, we performed sensory-specific satiation across 2 consecutive days, consisting of counterbalanced valued and devalued days. For the valued day, the mice were allowed to freely consume 20% sucrose solution for 1 hour. For the devalued day, mice were allowed to freely consume for 1 hour the pellet outcome previously earned in the lever press hold down task. One mouse that did not consume enough pellets (< 0.1 g) or sucrose (< 0.1 ml) during this free-access period was excluded from subsequent analysis (giving final n = 17, 10 males). Immediately following the feeding period, mice were placed into their respective operant chamber for a 10 minute session during which the number and duration of lever presses made were recorded, but no pellet reward was delivered. Investigators were not blind to the experimental groups. Response rate comparisons between valued and devalued days were made by normalizing each mouse’s test day response rate (RR) to the average response rate of their corresponding last 2 days of hold down training using the following formula:
*RRTest Day ÷ mean(RR1600ms4 + RR1600ms5)*

### Surgical Procedures

Mice first underwent isoflurane anesthesia (1-2%) before stereotaxic-guided intracranial injections via 500 nl volume Hamilton syringes (Reno, NV). Viral vectors were infused at a rate of 100 nl/minute and the syringe was then left unperturbed for 5 minutes to allow for diffusion after delivery. Mice were allowed to recover for a minimum of two weeks before the start of behavioral procedures. At the end of behavioral procedures, mice were euthanized and their brains were extracted and fixed in 4% paraformaldehyde. Optic fiber placement and viral expression was qualified by examining tracts in 50- to 100-μm-thick brain slices under a macro fluorescence microscope (Olympus MVX10). All surgical and behavioral experiments were performed during the light portion of the cycle.

For fiber photometry experiments, OFC was unilaterally targeted for viral injections at the following stereotaxic coordinates from Bregma: AP +2.7mm, L +1.65mm and V −2.6mm, with optic fiber ferrule placed V −2.5mm from the skull. For C57BL/6J mouse experiments, n = 9 mice (n = 6 males, n = 3 females) were injected with 300 nl of rAAV5/PAAV-CaMKIIa-GCaMP6s to express GCaMP6s under control of the Ca^2+^ calmodulin dependent protein kinase IIα (CamKIIα) promoter. For PV^cre^ mouse experiments, n = 8 mice (n = 5 males, n = 3 females) were injected with 300 nl of rAAV5/pAAV.CAG.Flex.GCaMP6s.WPRE.SV40 to express GCaMP6s via a Cre-dependent CAG promoter in PV+ neurons. An additional bilateral craniotomy was made over the posterior cerebellum for placement of screws to anchor a dental cement enclosure at the base of the ferrule to the base of the skull.

For optogenetic experiments, OFC was bilaterally targeted for viral injections at the following stereotaxic coordinates from Bregma: AP +2.6mm, L +1.75mm and V −2.1mm, with optic fiber ferrules placed V −1.9mm from the skull at a +12 degree orientation. For C57BL/6J mouse experiments, n = 12 mice (n = 7 males, n = 5 females) were injected with 250 nl of rAAV5/CamKII-hChR2(H134R)-eYFP-WPRE to express ChR2 under the CaMKIIα promoter for optogenetic activation or a combination of 250 nL of rAAV5/Ef1a-DIO-EYFP and 250 nl of rAAV5/CamKII-GFP-Cre for CamKIIα promoter fluorophore controls. For PV^cre^ mouse experiments, n = 14 mice (n = 10 males, n = 4 females) were injected with 250 nl of rAAV5/Ef1a-DIO-hChR2(H134R)-eYFP to express cre-dependent ChR2 in PV+ neurons for optogenetic activation or 250 nL of rAAV5/Ef1a-DIO-EYFP for fluorophore controls.

For lesion experiments, OFC was bilaterally targeted for viral injections at the following stereotaxic coordinates from Bregma: AP +2.7mm, L +1.65mm and V −2.6mm. C57BL/6J mice (n = 39, n = 27 males, n = 12 females) were injected with a combination of 250 nl of rAAV5/Ef1a-DIO-mCherry and 250 nL of rAAV5/AAV-Flex-taCasP3-TEVP for cre-dependent apoptosis lesions or 250 nl of rAAV5/Ef1a-DIO-mCherry for sham lesion controls. To assess the presence and spread of lesions, brains were first cut into 50 um slices and store at 4C in .1% sodium azide PBS before undergoing NeuN staining procedures using Alexa Fluor 488 Conjugate ABN78A4 Anti-NeuN (rabbit) antibody (Sigma-Aldrich). Slices were washed 3 times for 10 minutes with 1x PBS and pre-incubated in 10% Horse Serum and 0.3% Triton-X-100-PBS with 1% BSA for 1 hour. After, slices were incubated for 48 hours at 4C with primary antibody (1:500) in 2% horse serum and 0.3% Triton X-100-PBS-1% BSA 2%. Slices were then washed for 10 minutes with 3x PBS and stored at 4C until imaging.

### Fiber Photometry

After pre-training procedures, ferrule-implanted animals were unilaterally attached to bifurcated 400 um optical fiber tethers (Thorlabs, Newton, NJ) through which a 470nm LED (Thorlabs, Newton, NJ) excited virally expressed GCaMP6s (< 70 µW/mm2). Emitted fluorescence was monitored through the core of the bifurcated fiber using a 4x objective (Olympus, Shinjuku, Japan) focused onto a CMOS camera (FLIR Systems, Wilsonville, OR). Regions of interest demarcating each fiber fork were created within the fiber core using Bonsai software^105^ through which fluorescence intensity was captured at 20 Hz to produce two digitized signals, one for each animal connected to the bifurcated fiber. Analog behavioral timestamps for the beginning and end of each lever press, head entry, and reinforcer delivery periods were simultaneously sent to Bonsai software via TTL Med-PC pulses using microprocessors (Arduino Duo, from Arduino, Sumerville, MA) containing custom code. After each session, Bonsai software saved photometry signals and behavioral timestamps within comma-separated value files (.csv) that were then imported into Matlab (Mathworks Inc., Natick, MA) for subsequent analysis using custom scripts (see Code Availability). Raw fluorescence intensity signals underwent running median (5th order) and low pass (high cutoff frequency of 1 Hz) filtering to reduce noise and electrical artifacts. To correct for photobleaching in which a signal captured from fluorophores degrades by continuous light exposure during the session, we high pass filtered the signal with a low cutoff frequency of 0.001Hz. Filtered fluorescence intensity signals subsequently underwent a quality check for low expression and fiber decoupling. Briefly, sessions that did not exceed a 15 second moving window calculation of the signal’s 97.5 percentile by a minimum 1% fluorescence change^106^ or did not pass a visual inspection for within-session fiber-ferrule decoupling artifacts were excluded from further analyses. Peri-event changes in fluorescence intensity were then calculated via z-score normalization to each corresponding pre-lever press onset period (i.e. −5 seconds to −2 seconds prior to lever press). These z-scored fluorescence traces were then combined across all mice within a group to preserve the variance seen within a subject. Activity during the ongoing lever press duration was modified using Akima interpolation via MATLAB’s *interp1* function, excluding any lever press that was fewer than 2 samples (i.e. 100 ms) in duration that would invalidate interpolation. Comparisons between rewarded and unrewarded lever press traces were made using running permutation tests (1000 shuffles) that required at least 5 consecutive samples (or 3 consecutive samples for interpolated activity) to be different from one another.^77^ Population Ca^2+^ activity traces were then smoothed with MATLAB’s Savitzky–Golay *smoothdata* method using a 20 sample (or 1 sample for interpolated activity) sliding window for visual display purposes only.

### Optogenetic Excitation

Optogenetic excitation occurred only in the additional 5 sessions (days 6 to 10) during which the minimum duration criteria was 1600 ms. LEDs (470nm, Thorlabs) used for optogenetic excitation experiments were triggered by TTL pulses emitted from Med-PC operant chambers via Arduino Duos programmed with custom code. Sheathed (200 uM) optic fiber cables were coupled to bilaterally implanted ferrules (>= 1mW output at ferrule tip) through which a closed-loop system delivered light at 20 Hz (5 ms pulses) throughout the entirety of each (or every 7th) lever press duration.

### Linear Mixed Effects Models of Behavior

Linear Mixed Effects (LME) models were built to investigate the predictive relationship between the duration of individual lever presses (n) and the lever press occurring immediately prior to it (n - 1) ^33^. Random intercept terms for mouse and training day were included to account for the repeated, non-independent structure of the aggregated session data. To account for variance explained by the overall performance within a session, fixed terms included the overall percentage of rewarded lever presses as well as the timestamps of each lever press. To test how predictive relationships were contingent upon their sequential order, beta coefficient outputs pertaining to each behavioral measurement of interest were compared to a 1000 order shuffled (unless otherwise specified) distribution of beta coefficients using permutation testing (Table S1, top). Importantly, shuffling occurred within individual sessions/mice to preserve overall performance statistics (e.g. total lever presses made), and the order shuffling for each behavioral covariate occurred independently from each other. Thus the LME model for the behavioral cohort consisted of the following formula:

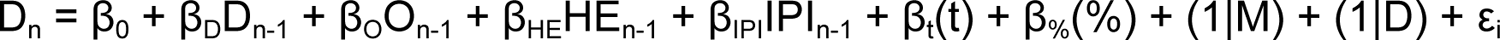

Where D_n_ is the current lever press duration, D_n-1_ is the prior lever press duration (in ms), O_n-1_ is the outcome of the prior lever press (binary 1 for reward, 0 for no reward), HE_n-1_ is the indicator of whether a head entry was made between the current and prior lever press (binary 1 for head entry made, 0 for no head entry made), IPI_n-1_ is the interpress interval (in ms), and B_x_ is the linear regression coefficient for each corresponding behavioral covariate term x (β_0_ being the intercept term). Covariates for lever press timestamps (t, in ms) and overall percentage of rewarded lever presses (%) were included alongside random intercept terms for mouse (M) and day (D).

To determine how far back the predictive relationship existed between press n and any particular n-back press, we built and 100 shuffled-order tested a similar LME model that included additional variables accounting for the duration of lever press n and n-back (n - 1 through n - 10) lever press durations as follows (Table S1, middle):

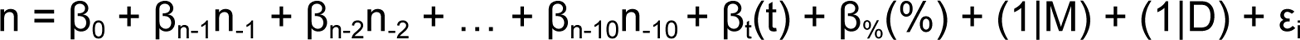

We built and 100 shuffled-order tested an additional LME model that included interaction terms to determine how prior behavioral variables (i.e. prior reward, checking, and interpress interval) compounded their effect on the subsequent lever press duration (Table S1, bottom):

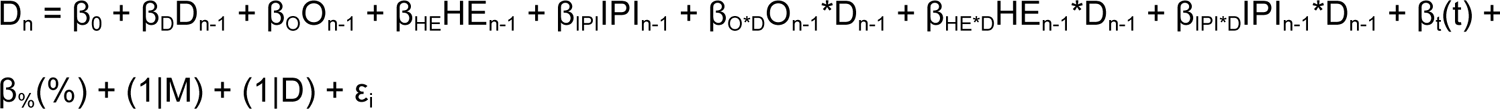

Regression coefficient terms β_x_ and shuffled-order testing procedures were as previously described, with the added covariates for main effects of prior duration (D_n-1_, lever press duration in ms) and its interactions with prior lever press outcome (O_n-1_*D_n-1_), prior presence of a head entry (HE_n-1_*D_n-1_), and interpress interval (IPI_n-1_*D_n-1_).

To determine how the predictive relationship of behavioral covariates for current lever press durations were affected by experimental manipulations (e.g. optogenetic excitation via ChR2), we built and 1000 shuffled-order tested similar LME models that included additional variables accounting for treatment group main effects and interactions as follows (Tables S4–S6):

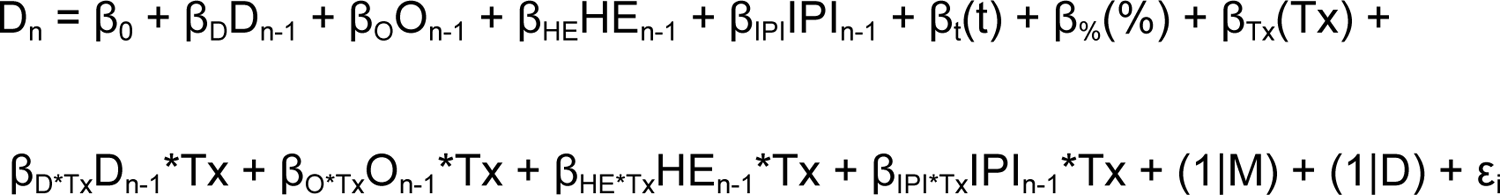

Regression coefficient terms β_x_ in these models were as previously described, with the added covariates for main effects of treatment (Tx, binary 1 for experimental and 0 for control groups) and its interactions with prior lever press duration (D_n-1_*Tx), prior outcome (O_n-1_*Tx), prior presence of a head entry (HE_n-1_*Tx), and interpress interval (IPI_n-1_*Tx). For the optogenetic excitation experiment in which only every 7th lever press triggered light delivery, a post-hoc LME model was tested using only the ChR2 group. In this case, however, the treatment main effect term Tx (and associated interaction terms) instead indicated the presence or absence of optogenetic stimulation for each individual lever press.

### Linear Mixed Effects Models of Ca^2+^ Activity

For OFC Ca^2+^ fluorescence activity monitoring experiments, LME models were built to predict Ca^2+^ activity within peri-event epochs given current and prior lever press durations alongside other behavioral variables. For these models, only data collected from the 1600 ms minimum duration criterion days were used. The mean area under the curve of activity traces during each of three epochs (−1s to 0s before lever press onset, lever press duration, and 0s to +1s after lever press release) was calculated to predict activity at each of these three time points using the following formula (Tables S2 and S3):

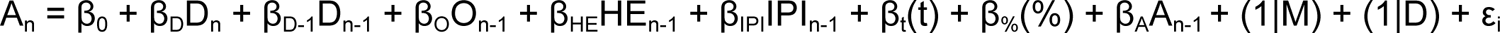

Where A_n_ is Ca^2+^ activity associated with the current lever press epoch (pre-onset, duration, or post-offset). Regression coefficient terms β_x_ in these models were as previously described, with the added covariates for main effects of current duration (D_n_, in ms) and prior lever press activity (A_n-1_) during that epoch to control for Ca^2+^ activity autocorrelation. LME regression coefficients for behavior measures of interest were compared to 1000 order shuffled datasets to test whether their predictive ability was due to the subsequent relationship.

### Quantification and Statistical Analysis

All analyses were two-tailed and statistical significance was defined as an α of p < 0.05. Statistical analysis was performed using GraphPad Prism 8.3.0 (GraphPad Software) and custom MATLAB R2019a (MathWorks) scripts using a PC desktop with Windows 10. Acquisition data, including lever presses, response rate, and percentage of lever presses that were rewarded were analyzed using one-way or two-way repeated measures ANOVAs with Greenhouse-Geisser corrections and Šidák corrections for post-hoc multiple comparisons unless otherwise noted. For outcome devaluation testing, a paired parametric t-test was performed to examine whether sensory-specific satiety reduced lever press responses on the devalued day compared to the valued day. For each LME model, we report the average regression coefficient (β), which measures the effect size and indicates how much a change in a predictor variable will change the output (e.g. lever press duration). Unless stated otherwise, significant predictors underwent follow-up permutation test comparisons for β coefficient values against a distribution of 1000 order or group shuffled versions of the same variable. For Ca^2+^ activity comparisons (i.e. reward vs no reward), permutation testing required 5 consecutive samples (or 3 consecutive samples for interpolated activity) that passed the threshold for significance. Lever presses longer than 10 seconds were excluded from all Ca^2+^ activity analyses. Data are presented as mean ± SEM.

## Key Resources Table

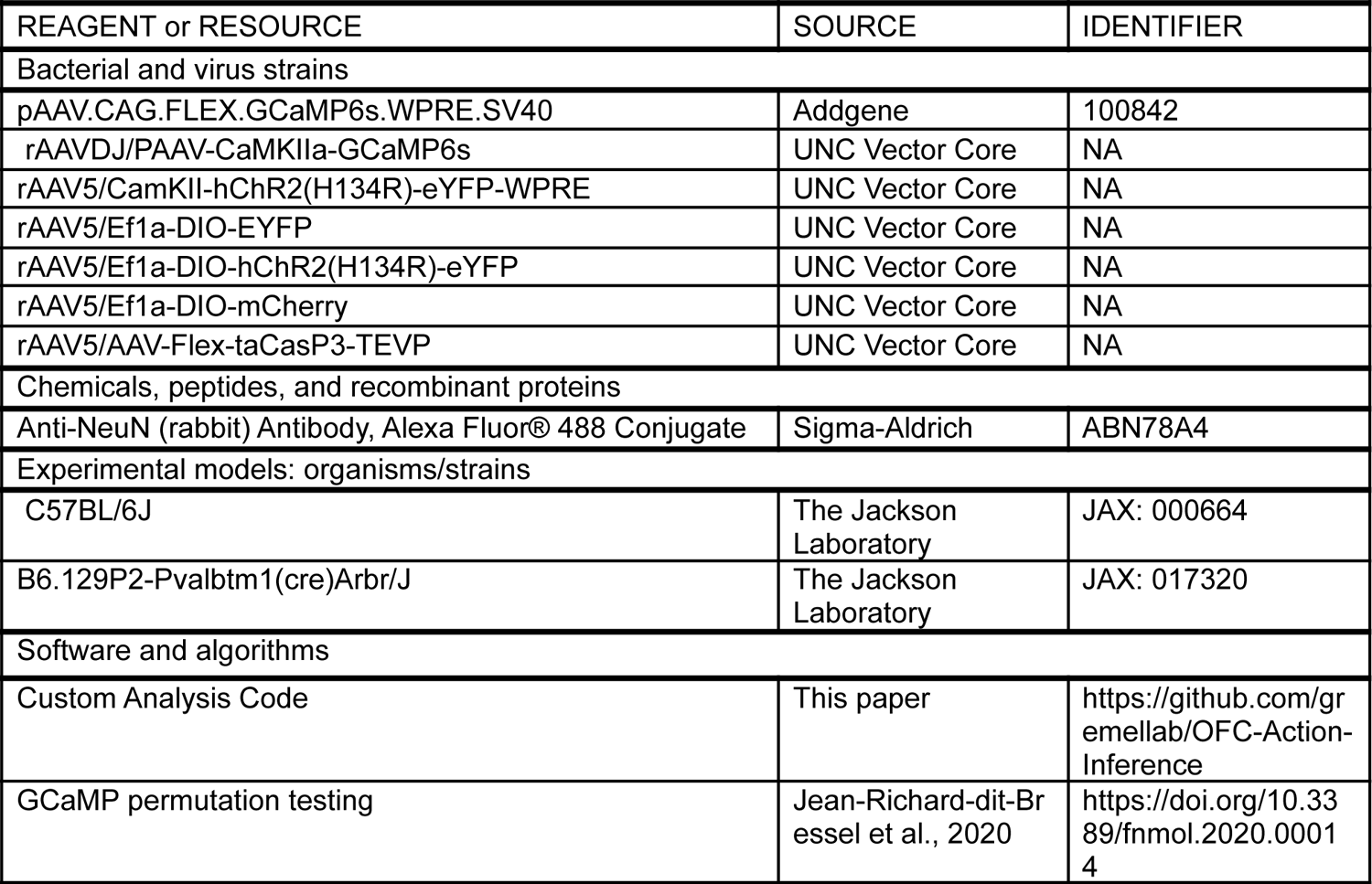

