## Supplementary Figures S1-4 and Tables S1-6 for "Orbitofrontal cortex populations are differentially recruited to support actions"

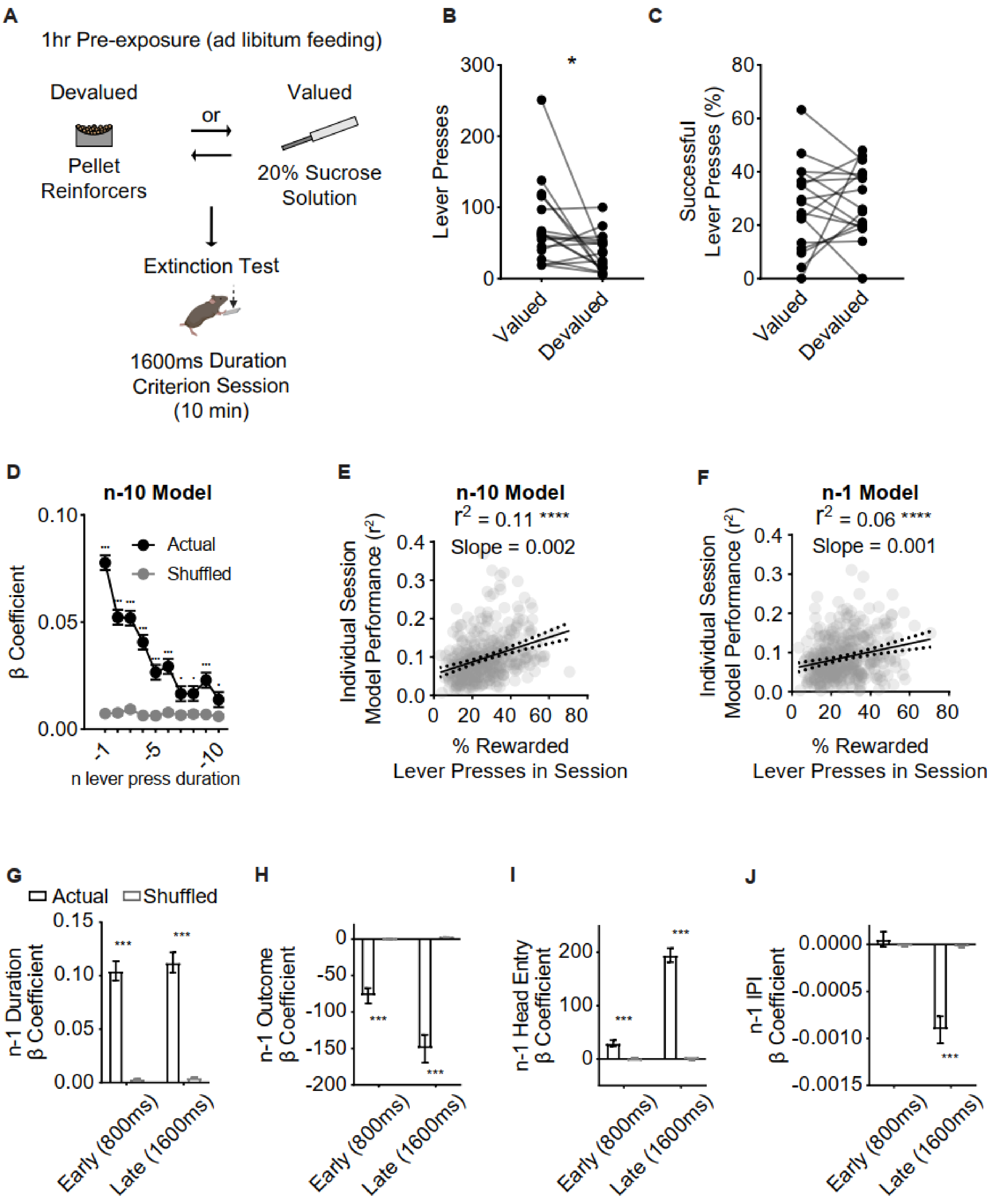


**Figure S1. Devaluation testing procedures, performance, and LME models relating prior lever press durations to current press duration and overall session performance, related to Figure 1.**

(A) Schematic showing counter-balanced devaluation testing procedures. Mice had 1 hour of free access to food pellets (devalued state) or a 20% sucrose solution (valued state) prior to a 10-minute 1600ms duration criterion session during which no reinforcers were delivered.

(B–C) Total lever presses (B) and percentage of lever presses (C) that exceeded the duration criterion in valued and devalued states throughout devaluation testing. Paired t tests revealed different patterns of lever pressing ($t_{16} = 2.408$, $p < 0.05$), but no difference in the percentage of lever presses ($t_{16} = 0.3536$, $p > 0.05$) that exceeded the duration criterion between valued and devalued states.

(D) β coefficients of LME model relating current lever press duration (n) to prior (n – 10) press durations for actual and order shuffled data.

(E) Linear regression for individual session data LME (n – 10) model described in (D), fitting ($R^2$) and percentage of lever presses that exceeded the duration criterion within that session. Dotted lines indicate 95% confidence interval of the best-fit line. Best-fit value slope = 0.001611 and Goodness of Fit ($R^2$) = 0.1120. Significant deviation from zero: $F_{1, 368} = 46.44$, $p < 0.0001$.

(F) Linear regression for individual session data LME (n – 1) model described in (Fig. 1K–N), fitting ($R^2$) and percentage of lever presses that exceeded the duration criterion within that session. Dotted lines indicate 95% confidence interval of the best-fit line. Best-fit value slope = 0.001062 and Goodness of Fit ($R^2$) = 0.05838. Significant deviation from zero: $F_{1, 368} = 22.81$, $p < 0.0001$.

(G–J) β coefficients of LME model relating current lever press duration (n) to prior (n – 1) press durations (G), press outcome (i.e. was lever press rewarded) (H), head entry (I), and interpress interval (IPI) (J) for actual and order shuffled data. Early 800 ms and Late 1600 ms refer to the first three days in which the duration criterion was >800 ms and the last two days in which the duration criterion >1600 ms, respectively.

Significance markers in (D) indicate comparisons to order shuffled data. Shuffled data are mean ± SEM of 1000 order shuffled β coefficients. * p < 0.05, *** p < 0.001, **** p < 0.0001.

See also Table S1 for more data.

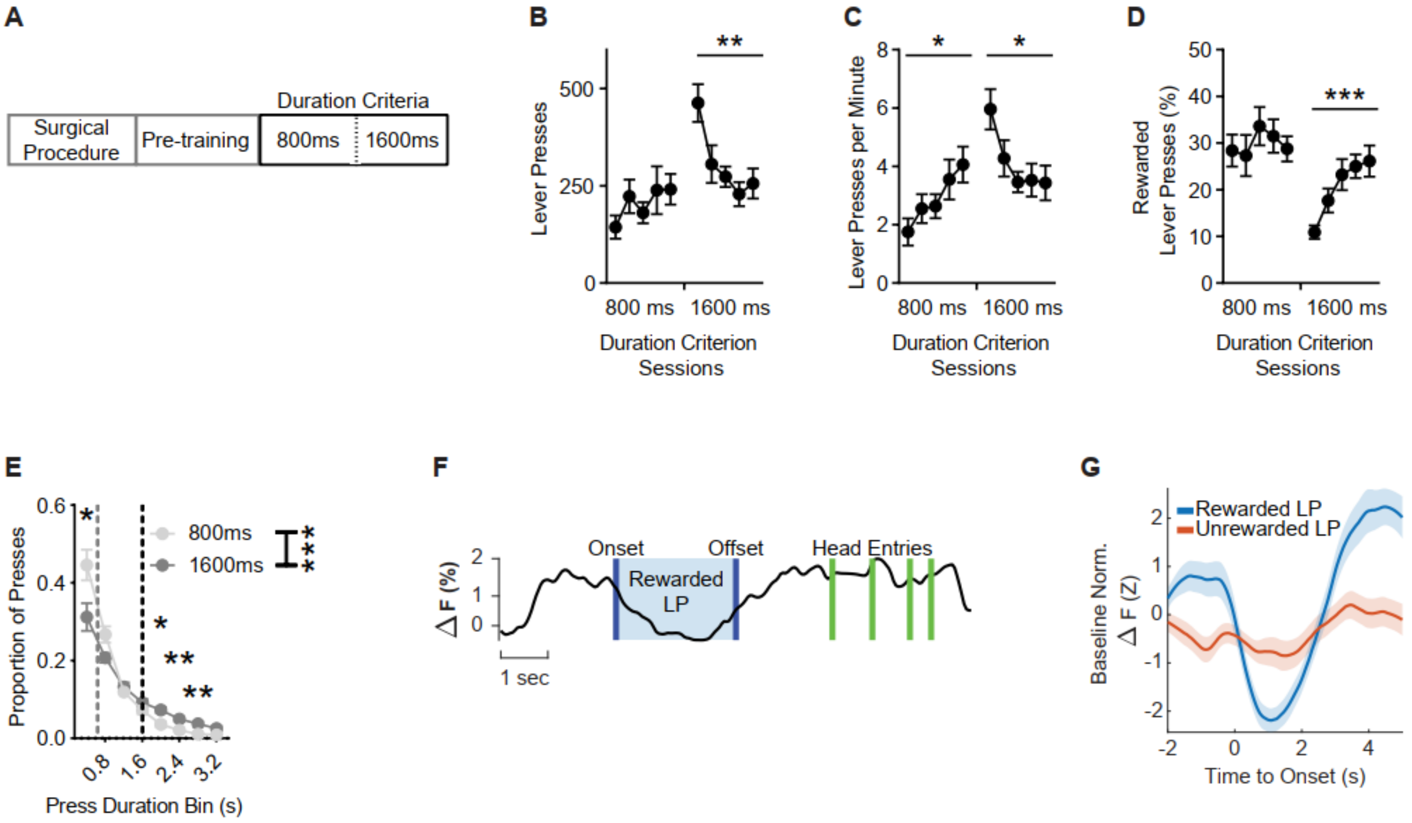


**Figure S2. Lever press performance throughout acquisition for fiber photometry experiments and representative OFC$^{CamKII+}$ $Ca^{2+}$ activity, related to Figure 2.**

(A) Training schedule for the lever press hold down task during fiber photometry experiments. Pretraining sessions were followed by sessions with a minimum duration criterion.

(B–D) Total lever presses (B), (C) lever pressing rate and (D) percentage of lever presses that exceeded the duration criterion across sessions. One-way mixed effects RM ANOVAs revealed a differences across 1600 ms sessions for total lever presses ($F_{2.589, 41.43} = 5.739$, $p < 0.01$), lever pressing rate ($F_{2.652, 42.43} = 3.712$, $p < 0.05$), and percentage of lever presses that exceeded the duration criterion ($F_{2.959, 47.35} = 7.535$, $p < 0.001$). A Mixed-effects analysis revealed a difference across 800 ms sessions for lever pressing rate only ($F_{2.650, 41.74} = 3.000$, $p < 0.05$).

(E) Histogram of lever press durations (400 ms bins) averaged for 800 ms and 1600 ms duration criterion sessions. A two-way mixed effects RM ANOVA (Bin * Criteria) on the proportion of presses across duration bins between the two duration criteria revealed a significant interaction: $F_{1.894, 30.30} = 9.954$, $p < 00.001$, a main effect of Bin: $F_{1.351, 21.61} = 95.29$, $p < 0.0001$, and a main effect of Criteria: $F_{1, 16} = 10.65$, $p < 0.01$). Post hoc comparisons showed significant differences between the duration criteria within the 0.4 s ($p < 0.05$), 2.0 s ($p < 0.05$), 2.4 s ($p < 0.01$), 2.8 s ($p < 0.01$) duration bins.

(F) Representative trace showing the percentage of changes in baseline normalized $Ca^{2+}$ activity over time. The region shaded in blue indicates a rewarded lever press duration. Green lines indicate head entries made.

(G) Representative $Ca^{2+}$ activity from a 1600 ms duration criterion session aligned to lever press onset (i.e.

initiation). Activity is z-score normalized to a pre-lever press onset baseline period. Blue indicates trace average of rewarded lever presses. Orange indicates trace average of unrewarded lever presses. LP = Lever Press. Data points represent mean ± SEM. * $p < 0.05$, ** $p < 0.01$, *** $p < 0.001$.

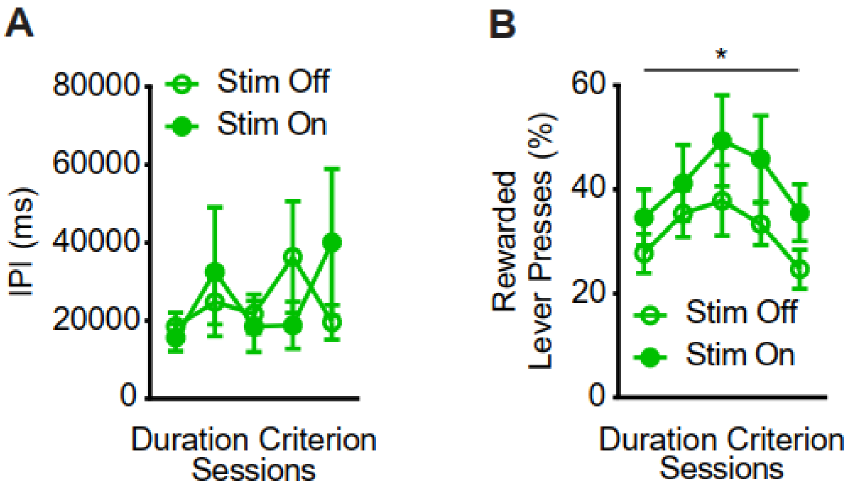


**Figure S3. Activation of OFC$^{CamKII+}$ neurons did not change the time interval to initiate subsequent lever press nor rewarded performance, related to Figure 4.**

(A) Interpress intervals from 1600 ms duration criterion sessions during which light was delivered on every 7th lever press, segmented by whether presses were paired with light activation or not (two-way RM ANOVA, no main effect of Stimulation $F_{1, 12} = 0.008$, $p = 0.93$ or Session $F_{4, 48} = 1.207$, $p = 0.40$).

(B) Percentage of lever presses that exceeded the duration criterion from 1600 ms duration criterion sessions during which light was delivered on every 7th lever press, segmented by whether presses were paired with light activation or not (two-way RM ANOVA, main effect of Session only $F_{4, 48} = 3.227$, $p = 0.02$).

Data points represent mean ± SEM. * $p < 0.05$

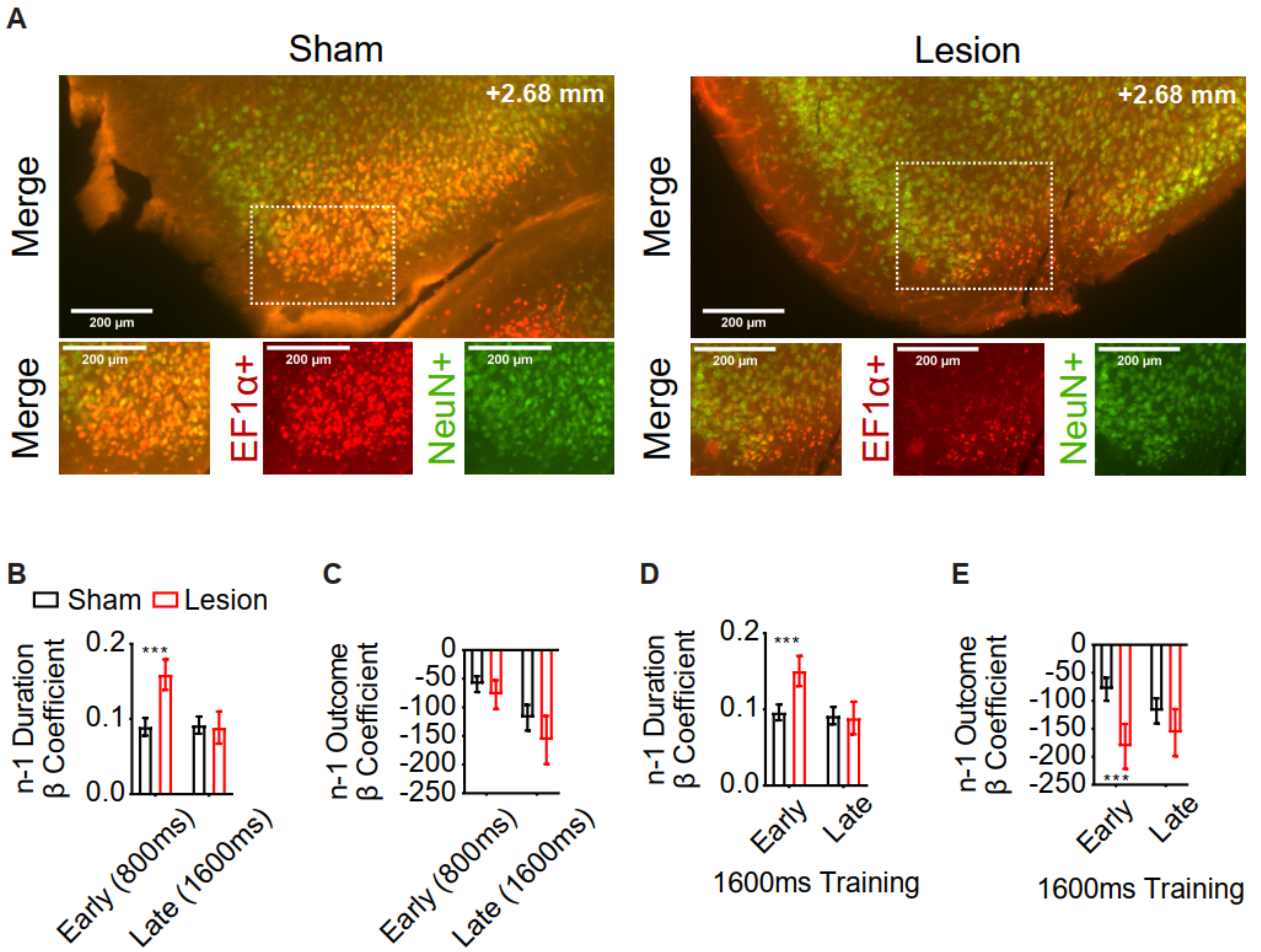


**Figure S4. Representative histology of sham and Cre-dependent caspase lesions of OFC$^{CamKII+}$ neurons and effects of lesions on model coefficients across training, related to Figure 5**.

(A) Representative histology as shown in (Figure 5A) of sham and Cre-dependent caspase lesions of lOFC neurons, zoomed in for visual clarity. Red indicates AAV-EF1α-DIO-mCherry expression. Green indicates immunohistochemical reactions for neural nuclear protein NeuN. White dotted squares indicate zoomed-in regions. Slice taken approximately from Bregma: AP +2.68mm, L +1.65mm and V −2.6mm.

(B and C) β coefficients of LME model relating current lever press duration (n) to prior (n - 1) press durations (B) and press outcome (i.e. was lever press rewarded) (C) for Sham and Lesion cohort actual data. Early 800 ms and Late 1600 ms refer to the first three days in which the duration criterion was >800 ms and the last two days in which the duration criterion was >1600 ms, respectively.

(D and E) β coefficients of LME model relating current lever press duration (n) to prior (n - 1) press durations (D) and press outcome (i.e. was lever press rewarded) (E) for Sham and Lesion cohort actual data. Early 1600 ms and Late 1600 ms refer to the first three days in which the duration criterion was >1600 ms and the last two days in which the duration criterion was >1600 ms, respectively.

Significance markers indicate comparisons to 1000 group shuffled data. Data points represent mean ± SEM. *** $p < 0.001$.

**Table S1. Behavior LME Model Statistics, related to Figure 1 and Figure S1.**

| n - 1 LME Model Statistics | | | | | | | |
|---|---|---|---|---|---|---|---|
| **Term** | **Coef** | **SE** | **Upper** | **Lower** | **F-Stat** | **F-Pval** | **Perm P** |
| Intercept | 314.8585 | 75.375065 | 462.59287 | 167.12412 | 17.449196 | 2.954E-05 | N/A |
| On-1 | -110.66545 | 7.4843838 | -95.996128 | -125.33476 | 218.63111 | 2.052E-49 | **p < 0.001** |
| Dn-1 | 0.1179562 | 0.0046091 | 0.12699 | 0.1089225 | 654.95565 | 6.04E-144 | **p < 0.001** |
| %Criteria | 14.042042 | 0.3167509 | 14.662871 | 13.421213 | 1965.2812 | 0 | **p < 0.001** |
| HEn-1 | 125.01432 | 4.8330364 | 134.48702 | 115.54162 | 669.08208 | 5.38E-147 | **p < 0.001** |
| Ts | 5.366E-05 | 1.612E-06 | 5.682E-05 | 5.05E-05 | 1108.161 | 1.57E-241 | **p < 0.001** |
| IPIn-1 | -0.0003987 | 5.602E-05 | -0.0002888 | -0.0005085 | 50.634357 | 1.121E-12 | **p < 0.001** |
| **n - 10 LME Model Statistics** | | | | | | | |
| **Term** | **Coef** | **SE** | **Upper** | **Lower** | **F-Stat** | **F-Pval** | **Perm P** |
| Intercept | 256.43136 | 57.342813 | 368.82276 | 144.03997 | 19.997895 | 7.762E-06 | N/A |
| Dn-1 | 0.077146 | 0.0033899 | 0.0837902 | 0.0705019 | 517.90671 | 2.59E-114 | **p < 0.01** |
| Dn-2 | 0.0516059 | 0.0034118 | 0.058293 | 0.0449188 | 228.7864 | 1.274E-51 | **p < 0.01** |
| Dn-3 | 0.0527951 | 0.0034231 | 0.0595043 | 0.0460858 | 237.87327 | 1.345E-53 | **p < 0.01** |
| Dn-4 | 0.0408967 | 0.0034323 | 0.047624 | 0.0341693 | 141.96957 | 1.046E-32 | **p < 0.01** |
| Dn-5 | 0.0259445 | 0.0034392 | 0.0326854 | 0.0192037 | 56.908314 | 4.61E-14 | **p < 0.01** |
| Dn-6 | 0.0293307 | 0.0034454 | 0.0360836 | 0.0225778 | 72.471728 | 1.721E-17 | **p < 0.01** |
| Dn-7 | 0.0165319 | 0.0034511 | 0.0232961 | 0.0097677 | 22.946572 | 1.668E-06 | **p = 0.03** |
| Dn-8 | 0.0160076 | 0.0034534 | 0.0227762 | 0.0092391 | 21.486694 | 3.568E-06 | **p = 0.01** |
| Dn-9 | 0.0233749 | 0.0034577 | 0.030152 | 0.0165978 | 45.700008 | 1.387E-11 | **p < 0.01** |
| Dn-10 | 0.0132799 | 0.0034536 | 0.0200489 | 0.0065108 | 14.785647 | 0.0001205 | **p = 0.02** |
| %Criteria | 10.679852 | 0.3384171 | 11.343147 | 10.016558 | 995.9236 | 2.28E-217 | **p < 0.01** |
| Ts | 3.288E-05 | 1.669E-06 | 3.615E-05 | 2.961E-05 | 388.16055 | 3.197E-86 | **p < 0.01** |
| **n - 1 Interactions LME Model Statistics** | | | | | | | |
| **Term** | **Coef** | **SE** | **Upper** | **Lower** | **F-Stat** | **F-Pval** | **Perm P** |
| Intercept | 335.25079 | 74.460159 | 481.19196 | 189.30963 | 20.271772 | 6.73E-06 | N/A |
| On-1 | -136.98086 | 11.867438 | -113.72081 | -160.24092 | 133.23119 | 8.45E-31 | N/A |
| Dn-1 | 0.0853273 | 0.0067159 | 0.0984903 | 0.0721643 | 161.42588 | 5.94E-37 | N/A |
| %Criteria | 14.10931 | 0.3168733 | 14.730378 | 13.488241 | 1982.6228 | 0 | N/A |
| HEn-1 | 51.356376 | 7.7364406 | 66.519722 | 36.19303 | 44.066276 | 3.19E-11 | N/A |
| Ts | 5.41E-05 | 1.61E-06 | 5.73E-05 | 5.09E-05 | 1127.7028 | 1.00E-245 | N/A |
| IPIn-1 | 5.57E-05 | 9.03E-05 | 0.0002326 | -0.0001213 | 0.3801208 | 0.5375401 | N/A |
| On-1 * Dn-1 | 0.0059397 | 0.0086302 | 0.0228548 | -0.0109754 | 0.473677 | 0.4913022 | **p = 0.69** |
| HEn-1 * Dn-1 | 0.0893464 | 0.0072528 | 0.1035619 | 0.075131 | 151.75403 | 7.64E-35 | **p < 0.01** |
| IPIn-1 * Dn-1 | -4.60E-07 | 6.69E-08 | -3.29E-07 | -5.91E-07 | 47.33278 | 6.03E-12 | **p < 0.01** |

Parameters, their coefficients, and statistical tests for LME models predicting n duration given prior behavior. Degrees of Freedom for all (top) F-tests = 1, 91286, (middle) F-tests = 1, 87950, (bottom) F-tests = 1, 91283. On-1 = Outcome of prior lever press as binary 1 for rewarded and 0 for unrewarded. Dn-x = Duration of x prior lever press in ms. %Criteria = Percentage of lever presses in session that exceeded duration criterion. HEn-1 =

Headentry between current and prior lever press as binary 1 for head entry present and 0 for head entry absent. Ts = Lever press session timestamp in ms. $IPI_{n-1}$ = time in ms between prior and current lever press. Data presented as regression coefficient variables that indicate their contribution in predicting current lever press duration changes given their occurrence. Data presented as two multiplied variables indicate the interaction with $D_{n-1}$ and shows how the contribution of prior lever press duration in predicting current lever press duration changes given that variable.

Coef = β Coefficient. SE = Standard Error. Upper and Lower = 95% confidence intervals. F-stat = F-statistic. F-Pval = p-value from the F-test. Perm P = p-value from permutation test comparing to 1000 order shuffled (top) or 100 order shuffled (middle and bottom) β coefficients. P-values of 0 are reported for some terms due to Matlab's numerical resolution.

**Table S2. OFC$^{CamKII+}$ Activity LME Model Statistics, related to Figure 2.**

| | Before Press, df = 15225 | | | |
|---|---|---|---|---|
| **Term** | **Coef** | **F** | **F-Pval** | **Perm P** |
| Intercept | -4.8539932 | 1.4322664 | 0.2314135 | N/A |
| Dn-0 | 0.0009882 | 3.9766795 | **0.0461522** | **p = 0.009** |
| Dn-1 | 6.914E-05 | 0.0341399 | 0.853412 | p = 0.846 |
| IPIn-1 | 6.418E-05 | 42.490124 | **7.327E-11** | N/A |
| On-1 | -12.525499 | 48.024991 | **4.377E-12** | N/A |
| HEn-1 | 19.514045 | 166.83156 | **5.763E-38** | N/A |
| Ts | 1.865E-06 | 24.689444 | **6.808E-07** | N/A |
| AUCn-1 | 0.3115896 | 1656.0424 | **0** | N/A |

| | During Press, df = 15146 | | | |
|---|---|---|---|---|
| **Term** | **Coef** | **F** | **F-Pval** | **Perm P** |
| Intercept | -1.426484 | 1.1669784 | 0.2800402 | N/A |
| Dn-0 | -1.06E-03 | 49.433901 | **2.14E-12** | **p < 0.001** |
| Dn-1 | 5.01E-04 | 19.450976 | **1.04E-05** | **p < 0.001** |
| IPIn-1 | 1.92E-05 | 41.2402 | **1.39E-10** | N/A |
| On-1 | -1.06E+00 | 3.7458827 | 0.0529563 | N/A |
| HEn-1 | 3.54E+00 | 59.665488 | **1.19E-14** | N/A |
| Ts | 7.80E-07 | 46.406142 | **9.97E-12** | N/A |
| AUCn-1 | 3.39E-01 | 1976.8657 | **0** | N/A |

| | After Press, df = 15225 | | | |
|---|---|---|---|---|
| **Term** | **Coef** | **F** | **F-Pval** | **Perm P** |
| Intercept | -2.5717668 | 0.1447539 | 0.7036053 | N/A |
| Dn-0 | 1.61E-02 | 95.102603 | **2.10E-22** | **p < 0.001** |
| Dn-1 | 8.33E-03 | 44.138628 | **3.16E-11** | **p < 0.001** |
| IPIn-1 | 4.08E-05 | 1.5410547 | 0.21448 | N/A |
| On-1 | -2.86E+01 | 22.480695 | **2.14E-06** | N/A |
| HEn-1 | -2.94E+01 | 33.454249 | **7.44E-09** | N/A |
| Ts | 4.48E-06 | 13.466452 | **0.0002437** | N/A |
| AUCn-1 | 3.84E-01 | 2652.6839 | **0** | N/A |

Parameters, their coefficients, and statistical tests relating CamKII+ OFC calcium activity to behavior. We predicted activity at three different epochs relative to the lever press: -1s to 0s before lever press initiation, during the lever press, and 0s to +5s after lever press termination. Dn-0 = Duration of current lever press in ms. Dn-1 = Duration of prior lever press in ms. On-1 = Outcome of prior lever press as binary 1 for rewarded and 0 for unrewarded. HEn-1 = Headentry between current and prior lever press as binary 1 for head entry present and 0 for head entry absent. Ts = Lever press session timestamp in ms. IPIn-1 = time in ms between prior and current lever press.

We included prior activity (AUCn-1) as a covariate to control for autocorrelation in calcium activity signal. Data presented as regression coefficient variables that indicate their contribution in predicting current activity changes given their occurrence. Bolded terms indicate significance in the model by F-test.

Coef = β Coefficient. F-Pval = p-value from the F-test. Perm P = p-value from permutation test comparing to 1000 order shuffled β coefficients. P-values of 0 are reported for some terms due to Matlab's numerical resolution.

**Table S3. OFC$^{PV+}$ Activity LME Model Statistics, related to Figure 2.**

| | Before Press, df = 9586 | | | |
|---|---|---|---|---|
| **Term** | **Coef** | **F** | **F-Pval** | **Perm P** |
| Intercept | -2.822237291 | 0.479099992 | 0.488847031 | N/A |
| Dn-0 | 6.58E-05 | 0.017171255 | 0.895747099 | p = 0.885 |
| Dn-1 | 0.00068703 | 1.96485418 | 0.1610275 | p = 0.155 |
| IPIn-1 | 8.04E-07 | 0.005635457 | 0.940160836 | N/A |
| On-1 | 4.346777874 | 4.985793376 | **0.025579259** | N/A |
| HEn-1 | 0.358177489 | 0.050383202 | 0.822402594 | N/A |
| Ts | -8.81E-08 | 0.046939823 | 0.82848105 | N/A |
| AUCn-1 | 0.433348986 | 2216.289164 | 0 | N/A |

| | During Press, df = 6972 | | | |
|---|---|---|---|---|
| **Term** | **Coef** | **F** | **F-Pval** | **Perm P** |
| Intercept | -1.254393297 | 0.734170862 | 0.391564031 | N/A |
| Dn-0 | 9.49E-05 | 0.305502025 | 5.80E-01 | p = 0.551 |
| Dn-1 | 6.99E-05 | 0.174332074 | 6.76E-01 | p = 0.639 |
| IPIn-1 | -2.56E-06 | 0.491696991 | 4.83E-01 | N/A |
| On-1 | 1.62E+00 | 5.938638458 | **0.014837301** | N/A |
| HEn-1 | 1.05E+00 | 3.563447132 | 5.91E-02 | N/A |
| Ts | 2.08E-08 | 0.015701231 | 9.00E-01 | N/A |
| AUCn-1 | 1.96E-01 | 278.0836691 | 2.98E-61 | N/A |

| | After Press, df = 9586 | | | |
|---|---|---|---|---|
| **Term** | **Coef** | **F** | **F-Pval** | **Perm P** |
| Intercept | 8.277871193 | 0.922054651 | 0.336960285 | N/A |
| Dn-0 | 8.29E-03 | 28.07694468 | **1.19E-07** | p = 0.691 |
| Dn-1 | 4.44E-04 | 0.084562547 | 7.71E-01 | **p < 0.001** |
| IPIn-1 | -3.03E-05 | 0.822718728 | 0.36440927 | N/A |
| On-1 | 3.37E+01 | 30.90719611 | **2.78E-08** | N/A |
| HEn-1 | -4.21E+01 | 70.43578476 | **5.43E-17** | N/A |
| Ts | 1.54E-06 | 1.481249398 | 0.223609106 | N/A |
| AUCn-1 | 3.65E-01 | 1441.454589 | **5.71E-294** | N/A |

Parameters, their coefficients, and statistical tests relating PV+ OFC calcium activity to behavior. We predicted activity at three different epochs relative to the lever press: -1s to 0s before lever press initiation, during the lever press, and 0s to +5s after lever press termination. Dn-0 = Duration of current lever press in ms. Dn-1 = Duration of prior lever press in ms. On-1 = Outcome of prior lever press as binary 1 for rewarded and 0 for unrewarded. HEn-1 = Headentry between current and prior lever press as binary 1 for head entry present and 0 for head entry absent. Ts = Lever press session timestamp in ms. IPIn-1 = time in ms between prior and current lever press.

We included prior activity (AUCn-1) as a covariate to control for autocorrelation in calcium activity signal. Data presented as regression coefficient variables that indicate their contribution in predicting current activity changes given their occurrence. Bolded terms indicate significance in the model by F-test.
Coef = β Coefficient. F-Pval = p-value from the F-test. Perm P = p-value from permutation test comparing to 1000 order shuffled β coefficients.

**Table S4. $OFC^{PV+}$ Activation Treatment Group LME Model Statistics, related to Figure 3.**

| Term | Coef | SE | Upper | Lower | F-Stat | F-Pval | Perm P |
|---|---|---|---|---|---|---|---|
| Intercept | 294.80624 | 66.96325 | 426.06986 | 163.54261 | 19.38208 | **1.08E-05** | N/A |
| On-1 | -158.34336 | 68.28033 | -24.49795 | -292.18878 | 5.37785 | **2.04E-02** | N/A |
| Dn-1 | 0.08201 | 0.02407 | 0.12919 | 0.03483 | 11.60913 | **6.59E-04** | N/A |
| %Criteria | 26.34108 | 1.99946 | 30.26050 | 22.42166 | 173.55614 | **2.91E-39** | N/A |
| HEn-1 | -43.47924 | 51.17692 | 56.83949 | -143.79797 | 0.72180 | **3.96E-01** | N/A |
| Ts | 0.00005 | 0.00001 | 0.00007 | 0.00004 | 45.91469 | **1.31E-11** | N/A |
| IPIn-1 | 0.00016 | 0.00021 | 0.00058 | -0.00026 | 0.55811 | 0.4550441 | N/A |
| Treatment | 38.18195 | 53.53097 | 143.11517 | -66.75127 | 0.50875 | 0.4756986 | N/A |
| On-1 * Treatment | 13.69072 | 91.86767 | 193.77284 | -166.39139 | 0.02221 | 0.8815361 | p = 0.803 |
| Dn-1 * Treatment | -0.07428 | 0.03290 | -0.00979 | -0.13878 | 5.09802 | **0.0239776** | **p = 0.006** |
| HEn-1 * Treatment | 130.74381 | 68.35552 | 264.73661 | -3.24900 | 3.65843 | 0.0558185 | **p = 0.01** |
| IPIn-1 * Treatment | -0.00043 | 0.00030 | 0.00017 | -0.00102 | 1.95460 | 0.1621277 | p = 0.118 |

Parameters, their coefficients, and statistical tests for LME model predicting n duration given prior behavior. Degrees of Freedom for all F-tests = 1, 8793. On-1 = Outcome of prior lever press as binary 1 for rewarded and 0 for unrewarded. Dn-1 = Duration of prior lever press in ms. %Criteria = Percentage of lever presses in session that exceeded duration criterion. HEn-1 = Headentry between current and prior lever press as binary 1 for head entry present and 0 for head entry absent. Ts = Lever press session timestamp in ms. IPIn-1 = time in ms between prior and current lever press. Treatment = Presence of excitatory opsin ChR2 as binary 1 for present and 0 for absent.

Data presented as two multiplied variables indicate the effect if they occur in ChR2 mice (e.g., the interaction between Dn-1 * Treatment shows how the contribution of prior lever press duration in predicting current lever press duration changes given presence of ChR2.

Coef = β Coefficient. SE = Standard Error. Upper and Lower = 95% confidence intervals. F-stat = F-statistic. F-Pval = p-value from the F-test. Perm P = p-value from permutation test comparing to 1000 order shuffled β coefficients.

**Table S5. OFC$^{CamKII+}$ Activation Treatment Group LME Model Statistics, related to Figure 4.**

| n -1 Treatment Interaction LME Model Statistics | | | | | | | |
|---|---|---|---|---|---|---|---|
| **Term** | **Coef** | **SE** | **Upper** | **Lower** | **F-Stat** | **F-Pval** | **Perm P** |
| Intercept | 371.7199 | 65.0241 | 499.1803 | 244.2595 | 32.6801 | **1.12E-08** | N/A |
| On-1 | -68.4277 | 71.8824 | 72.4764 | -209.3318 | 0.9062 | **3.41E-01** | N/A |
| Dn-1 | 0.0751 | 0.0231 | 0.1204 | 0.0297 | 10.5282 | **1.18E-03** | N/A |
| %Criteria | 26.5936 | 1.4276 | 29.3920 | 23.7952 | 347.0053 | **3.74E-76** | N/A |
| HEn-1 | -54.2107 | 53.5572 | 50.7723 | -159.1938 | 1.0246 | **3.11E-01** | N/A |
| Ts | 0.0001 | 0.0000 | 0.0001 | 0.0000 | 46.3795 | **1.03E-11** | N/A |
| IPIn-1 | -0.0003 | 0.0002 | 0.0001 | -0.0006 | 1.8466 | 0.174204 | N/A |
| Treatment | -139.1133 | 63.6851 | -14.2776 | -263.9490 | 4.7716 | **0.028957** | N/A |
| On-1 * Treatment | -171.2101 | 88.1492 | 1.5802 | -344.0004 | 3.7724 | **0.052132** | **p = 0.001** |
| Dn-1 * Treatment | 0.0863 | 0.0299 | 0.1449 | 0.0277 | 8.3388 | **0.003889** | **p = 0.001** |
| HEn-1 * Treatment | 213.2290 | 67.3024 | 345.1554 | 81.3025 | 10.0376 | **0.001538** | **p < 0.001** |
| IPIn-1 * Treatment | 0.0000 | 0.0003 | 0.0006 | -0.0006 | 0.0002 | 0.988174 | p = 0.989 |
| **ChR2 Mice Only: n -1 Activation Interaction LME Model Statistics** | | | | | | | |
| **Term** | **Coef** | **SE** | **Upper** | **Lower** | **F-Stat** | **F-Pval** | **Perm P** |
| Intercept | 164.6687 | 60.49933 | 283.2681 | 46.06925 | 7.408338 | **0.00651** | N/A |
| On-1 | -371.315 | 71.01612 | -232.099 | -510.531 | 27.33825 | **1.76E-07** | N/A |
| Dn-1 | 0.124286 | 0.033771 | 0.19049 | 0.058082 | 13.54388 | **0.000235** | N/A |
| %Criteria | 27.05511 | 1.525669 | 30.04594 | 24.06428 | 314.469 | **1.08E-68** | N/A |
| HEn-1 | 153.9502 | 63.7735 | 278.9682 | 28.93232 | 5.827472 | **0.015806** | N/A |
| Ts | 9.94E-05 | 1.27E-05 | 0.000124 | 7.46E-05 | 61.69031 | **4.7E-15** | N/A |
| IPIn-1 | 0.000224 | 0.000419 | 0.001044 | -0.0006 | 0.285319 | 0.593255 | N/A |
| Stimn-0 | 253.9721 | 58.90581 | 369.4477 | 138.4965 | 18.58899 | **1.65E-05** | N/A |
| Stimn-1 | 23.33592 | 60.98068 | 142.8789 | -96.2071 | 0.146442 | 0.701972 | N/A |
| Dn-1 * On-1 | 0.092325 | 0.039999 | 0.170738 | 0.013913 | 5.327664 | **0.021022** | N/A |
| Dn-1 * HEn-1 | 0.013083 | 0.032578 | 0.076947 | -0.05078 | 0.16127 | 0.688004 | N/A |
| Dn-1 * IPIn-1 | -5.4E-07 | 2.06E-07 | -1.4E-07 | -9.5E-07 | 6.974631 | **0.008288** | N/A |
| Dn-1 * Stimn-0 | -0.00982 | 0.034645 | 0.058093 | -0.07774 | 0.080375 | 0.7768 | N/A |
| IPIn-1 * Stimn-0 | 0.001341 | 0.000744 | 0.0028 | -0.00012 | 3.245466 | 0.071669 | N/A |
| Dn-1 * Stimn-1 | -0.07602 | 0.031821 | -0.01364 | -0.1384 | 5.706848 | **0.016928** | N/A |
| IPIn-1 * Stimn-1 | -0.00026 | 0.000634 | 0.000982 | -0.0015 | 0.168666 | 0.681314 | N/A |
| **YFP Mice Only: n -1 Activation Interaction LME Model Statistics** | | | | | | | |
| **Term** | **Coef** | **SE** | **Upper** | **Lower** | **F-Stat** | **F-Pval** | **Perm P** |
| '(Intercept)' | 483.75934 | 114.51877 | 708.28646 | 259.23223 | 17.844541 | **2.46E-05** | N/A |
| On-1 | -68.640362 | 107.18101 | 141.50024 | -278.78097 | 0.4101318 | 0.5219428 | N/A |
| Dn-1 | 0.0862542 | 0.0533197 | 0.1907935 | -0.0182852 | 2.6168871 | 0.1058181 | N/A |
| %Criteria | 23.272496 | 3.0809257 | 29.313002 | 17.231989 | 57.058909 | **5.31E-14** | N/A |
| HEn-1 | 54.43965 | 88.774153 | 228.49151 | -119.61221 | 0.3760604 | 0.5397574 | N/A |
| Ts | 2.73E-05 | 1.43E-05 | 5.52E-05 | -6.55E-07 | 3.6660641 | 0.0556093 | N/A |

| | | | | | | | |
|---|---|---|---|---|---|---|---|
| IPIn-1 | -3.35E-05 | 0.000382 | 0.0007155 | -0.0007824 | 0.0076729 | 0.9302032 | N/A |
| Stimn-0 | 164.82679 | 84.8703 | 331.2247 | -1.5711238 | 3.7717605 | 0.0522016 | N/A |
| Stimn-1 | 120.283 | 87.109118 | 291.07037 | -50.504366 | 1.9066952 | 0.1674154 | N/A |
| Dn-1 * On-1 | 0.0260959 | 0.0606194 | 0.1449473 | -0.0927554 | 0.18532 | 0.6668661 | N/A |
| Dn-1 * HEn-1 | -0.0676451 | 0.041791 | 0.014291 | -0.1495811 | 2.620033 | 0.1056086 | N/A |
| Dn-1 * IPIn-1 | -1.44E-07 | 2.39E-07 | 3.24E-07 | -6.13E-07 | 0.3641353 | 0.546256 | N/A |
| Dn-1 * Stimn-0 | -0.0216541 | 0.0494551 | 0.0753083 | -0.1186164 | 0.1917152 | 0.6615188 | N/A |
| IPIn-1 * Stimn-0 | -8.17E-05 | 0.0004492 | 0.0007991 | -0.0009625 | 0.0331018 | 0.8556407 | N/A |
| Dn-1 * Stimn-1 | -0.0887973 | 0.0465705 | 0.0025094 | -0.180104 | 3.6356161 | 0.0566341 | N/A |
| IPIn-1 * Stimn-1 | 6.10E-05 | 0.0007855 | 0.0016011 | -0.0014791 | 0.00603 | 0.9381083 | N/A |

Parameters, their coefficients, and statistical tests for LME models predicting n duration given prior behavior. Degrees of Freedom for all (top) F-tests = 1, 9936 (middle) F-tests = 1, 6266, (bottom) F-tests = 1, 3650. $O_{n-1}$ = Outcome of prior lever press as binary 1 for rewarded and 0 for unrewarded. $D_{n-1}$ = Duration of prior lever press in ms. %Criteria = Percentage of lever presses in session that exceeded duration criterion. $HE_{n-1}$ = Headentry between current and prior lever press as binary 1 for head entry present and 0 for head entry absent. Ts = Lever press session timestamp in ms. $IPI_{n-1}$ = time in ms between prior and current lever press. Treatment = Presence of excitatory opsin ChR2 as binary 1 for present and 0 for absent. Stimn-0 = Light stimulation during current lever press as binary 1 for light-on and 0 for light-off. Stimn-1 = Light stimulation during prior lever press as binary 1 for light-on and 0 for light-off.

For Treatment LME model (top), data presented as two multiplied variables indicate the effect if they occur in ChR2 mice (e.g., the interaction between $D_{n-1}$ * Treatment shows how the contribution of prior lever press duration in predicting current lever press duration changes given presence of ChR2.

For Stimulation LME models (middle and bottom), data presented as two multiplied variables also indicate the effect if they occur in current (n-0) or prior (n-1) lever presses (e.g., the interaction between Dn-1 * Stimn-0 shows how the contribution of prior lever press duration in predicting current lever press duration changes given presence of light stimulation.

Coef = β Coefficient. SE = Standard Error. Upper and Lower = 95% confidence intervals. F-stat = F-statistic. F-Pval = p-value from the F-test. Perm P = p-value from permutation test comparing to 1000 order shuffled β coefficients.

**Table S6. OFC$^{CamKII+}$ Lesion Treatment Group LME Model Statistics, related to Figure 5.**

| Term | Coef | SE | Upper | Lower | F-Stat | F-Pval | Perm P |
|---|---|---|---|---|---|---|---|
| '(Intercept)' | 331.88952 | 69.65927 | 468.42071 | 195.35833 | 22.70018 | **1.90E-06** | N/A |
| On-1 | -86.11355 | 8.92831 | -68.61418 | -103.61292 | 93.02594 | **5.27E-22** | N/A |
| Dn-1 | 0.10801 | 0.00531 | 0.11841 | 0.09761 | 414.43529 | **5.91E-92** | N/A |
| %Criteria | 14.03897 | 0.25970 | 14.54798 | 13.52996 | 2922.27079 | **0.00E+00** | N/A |
| HEn-1 | 81.63634 | 5.57521 | 92.56368 | 70.70900 | 214.40969 | **1.67E-48** | N/A |
| Ts | 0.00004 | 0.00000 | 0.00005 | 0.00004 | 887.46022 | **3.21E-194** | N/A |
| IPIn-1 | 0.00000 | 0.00006 | 0.00012 | -0.00013 | 0.00370 | 0.9515054 | N/A |
| Treatment | 12.87046 | 17.96350 | 48.07867 | -22.33775 | 0.51334 | 0.4736976 | N/A |
| On-1 * Treatment | -47.03541 | 13.57523 | -20.42815 | -73.64268 | 12.00481 | **0.0005308** | **p < 0.001** |
| Dn-1 * Treatment | 0.03874 | 0.00758 | 0.05360 | 0.02388 | 26.12031 | **3.21E-07** | **p < 0.001** |
| HEn-1 * Treatment | -4.66682 | 9.16477 | 13.29601 | -22.62964 | 0.25930 | 0.6106043 | p = 0576 |
| IPIn-1 * Treatment | 0.00008 | 0.00010 | 0.00027 | -0.00011 | 0.65884 | 0.4169703 | p = 0.446 |

Parameters, their coefficients, and statistical tests for LME model predicting n duration given prior behavior. Degrees of Freedom for all F-tests = 1, 108014. On-1 = Outcome of prior lever press as binary 1 for rewarded and 0 for unrewarded. Dn-1 = Duration of prior lever press in ms. %Criteria = Percentage of lever presses in session that exceeded duration criterion. HEn-1 = Headentry between current and prior lever press as binary 1 for head entry present and 0 for head entry absent. Ts = Lever press session timestamp in ms. IPIn-1 = time in ms between prior and current lever press. Treatment = Presence of caspase Lesion as binary 1 for present and 0 for absent.

Data presented as two multiplied variables indicate the effect if they occur in Lesioned mice (e.g., the interaction between Dn-1 * Treatment shows how the contribution of prior lever press duration in predicting current lever press duration changes given presence of caspase Lesion.

Coef = β Coefficient. SE = Standard Error. Upper and Lower = 95% confidence intervals. F-stat = F-statistic. F-Pval = p-value from the F-test. Perm P = p-value from permutation test comparing to 1000 order shuffled β coefficients.
